# Metacognition facilitates the exploitation of unconscious brain states

**DOI:** 10.1101/548941

**Authors:** Aurelio Cortese, Hakwan Lau, Mitsuo Kawato

## Abstract

Can humans be trained to make strategic use of unconscious representations in their own brains? We investigated how one can derive reward-maximizing choices from latent high-dimensional information represented stochastically in neural activity. In a novel decision-making task, reinforcement learning contingencies were defined in real-time by fMRI multivoxel pattern analysis; optimal action policies thereby depended on multidimensional brain activity that took place below the threshold of consciousness. We found that subjects could solve the task, when their reinforcement learning processes were boosted by implicit metacognition to estimate the relevant brain states. With these results we identified a frontal-striatal mechanism by which the brain can untangle tasks of great dimensionality, and can do so much more flexibly than current artificial intelligence.

---

We consciously perceive our reality, yet much of ongoing brain activity is unconscious (*1, 2*). While such activity may contribute to behaviour, presumably it does so automatically and is not strategically utilized. Can humans be trained to make rational use of this rich, unconscious brain activity? From the outset, this problem is challenging because the relevant activity is often high dimensional; given so many *unconscious* dimensions, how can subjects know what to learn?

Previous studies have shown that reinforcement learning (RL) can operate on *external* masked stimuli (*3*). In those studies, the relevant subliminal information was driven by a simple visual stimulus, which carried only a single bit of information. Here we address a somewhat more challenging question with a novel technique based on internal multivariate representations. Subjects have to learn a ‘hidden’ brain state with many dimensions generated stochastically within the brain.

Using a machine learning classifier (a ‘decoder’), subjects’ brain activity (in prefrontal cortex – PFC, or visual cortex – VC^1^) determined in real-time the reward contingencies in a RL task (see figure 1, supplementary methods). Briefly, a decoder classified motion direction even though the stimulus presented had no coherent motion – essentially classifying stochastic brain activity based on its resemblance to left or right motion. While no motion information was presented, subjects had to make a perceptual discrimination (report rightward or leftward motion direction), and then rate their confidence in their choice. Afterwards, they had to gamble on two options (A or B) that could potentially lead to reward (30 ¥). Unbeknownst to the subjects, whether it was option A or B that was more likely to be rewarded (i.e. the optimal action) was determined by a multidimensional pattern of brain activity, that are known to be unconscious (*1, 4, 5*). That is, the decoded motion direction was used to determine reward contingencies.

**Figure 1:**
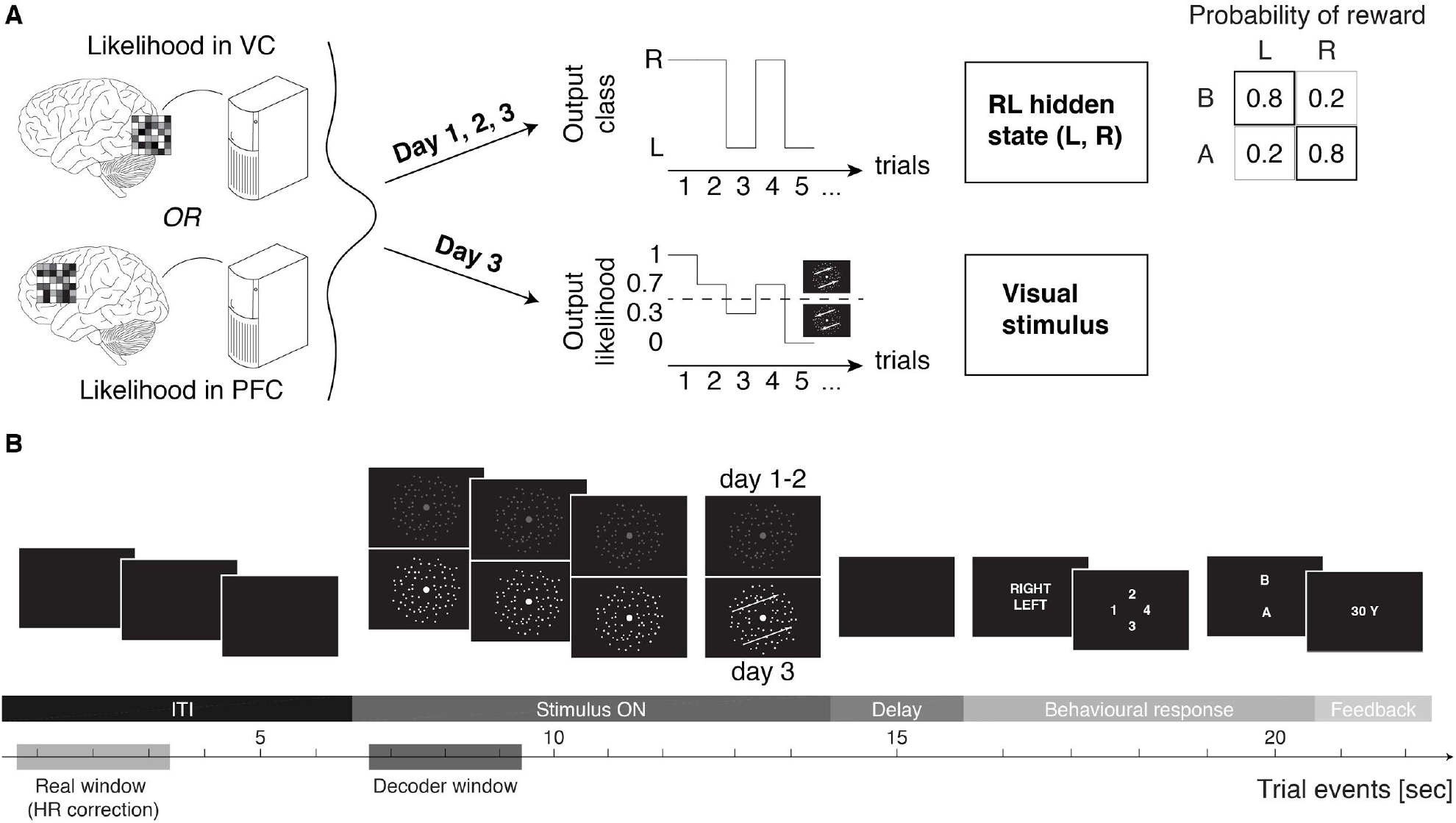
Design of the unconscious-brain-state reinforcement learning task. Prior to the learning task subjects engaged in a simple dot motion discrimination task in order to acquire fMRI data to construct their individual motion representation decoders (see supplementary figure 1 and methods). **A**, the learning task consisted of 3 consecutive days. On each day, decoding was performed with fMRI multivoxel patterns from either the visual cortex (VC) or prefrontal cortex (PFC), depending on the group to which a subject was assigned. On all days, the decoder output was used in real-time to determine the RL states “(i.e., unconscious representation of leftward or rightward motion)” on a trial-by-trial basis. In a given state, only one action was optimal, with high probability (0.8) of reward, while the other action had low reward probability (0.2). On the last day the decoder output likelihood was also used to proportionally define the motion direction (see supplementary methods). **B**, each trial started with a blank inter-trial interval (ITI, 6 sec). Random dot motion was then shown for 8 sec (Stimulus ON). On the first two days, the motion was entirely random, while on the 3rd day the last 2 sec of Stimulus ON had increasingly higher coherence, as determined by the decoder output likelihood. After a 1 sec delay, subjects had to report the direction of motion (the unconscious state), their confidence on the visual discrimination, and then gamble on one of two actions (A or B). Following action selection, the outcome for the current trial (reward: 30¥ [0.25$], or no reward: 0¥/$) was shown on the screen. Decoding was performed with 3 data points starting from the first TR of stimulus ON, and the three output likelihoods were then averaged to a single value. Because of the hemodynamic delay, this effectively amounted to using brain activity during the first 3 TRs of the ITI, ruling out possible confounds due to random dot motion viewing. The final averaged likelihood was input into a Heaviside step function to define the label of the current trial and therefore the contingency for the action-reward rule: *Label = L for likel < 0.5 Label = R for likel* > 0.5. HR: hemodynamic response delay, L: left, R: right.

Given the unconscious nature of the critically relevant information, it may seem improbable that subjects can learn to perform advantageously in this task. However, previously we have proposed that such problems may be solved via the mechanism of metacognition (*6, 7*). We hypothesized that by creating low-dimensional representations in the prefrontal cortex (similar to the ‘chunking’ phenomenon in working memory (*8*)), metacognition may accelerate RL because operations occur in a reduced state-space rather than the original, multidimensional one. This interaction between RL and metacognition could depend on parallel loops linking frontal and striatal brain regions.

As anticipated, we found that subjects could indeed learn to perform well in the gambling task, confirming our prediction.

## Learning to use unconscious information, behavioural and computational accounts

Over the course of approximately two hundred trials, subjects showed evidence of learning, with above chance reward-maximizing action selection (all results report mean probability ± s.e.m., one-sided t-test, chance level 0.5; day 1: 0.524 ± 0.016 [t_17_ = 1.49, *P* = 0.077], day 2: 0.537 ± 0.012 [t_17_ = 3.25, *P* = 0.0024], figure 2A). This happened despite the fact that the unconscious brain states relevant for selecting the optimal action were not physically presented to the subjects, and that their perceptual discrimination was no better than chance (figure 2B). If subjects were actually conscious about the brain state taken as the output of the decoder, then this information should have been used for the perceptual decision and discrimination accuracy would have been better than chance.

**Figure 2:**
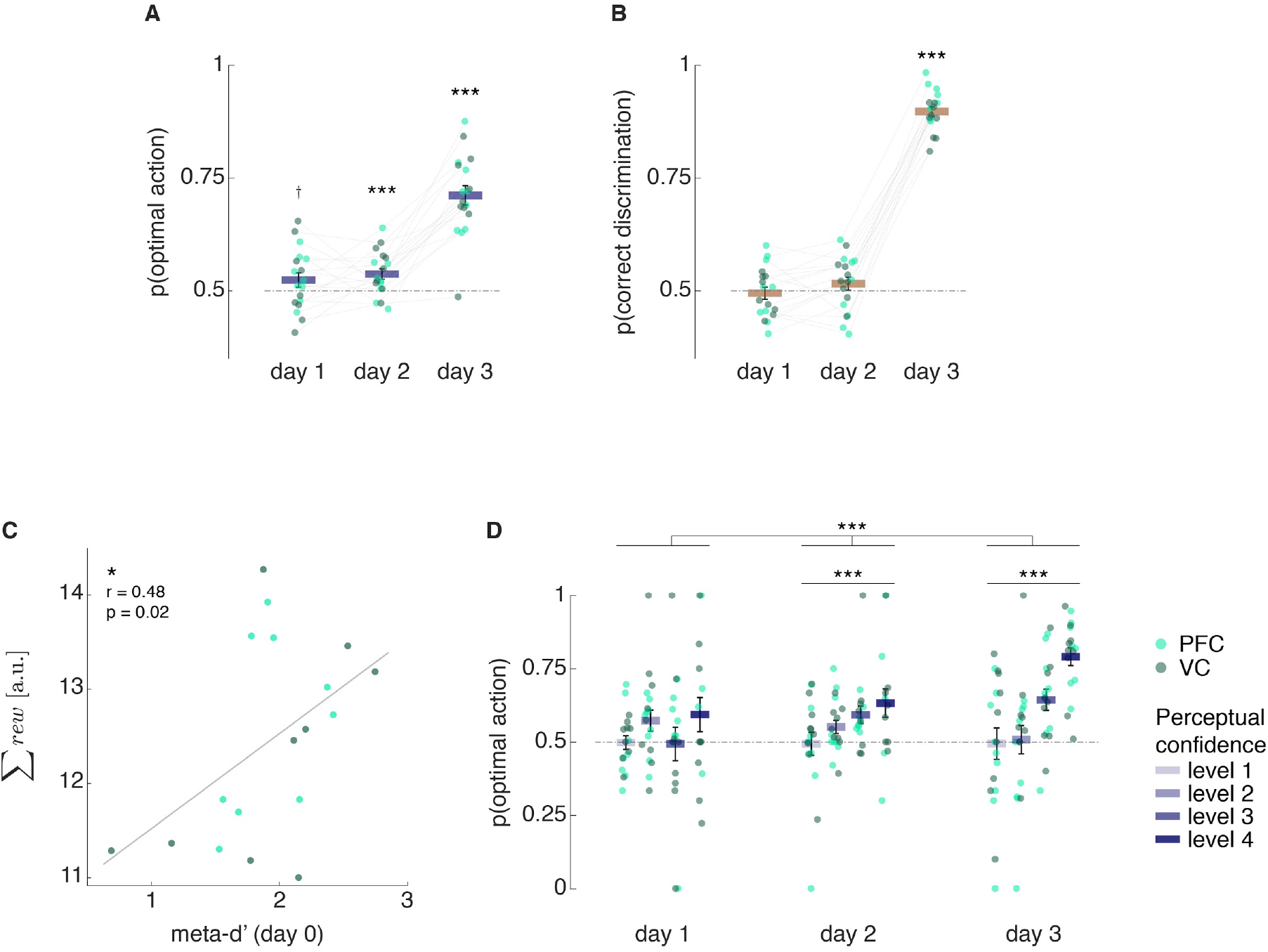
Analyses of behaviour: learning to use unconscious brain states and the contribution of metacognition. Subjects learned to associate unconscious brain states with specific actions that were more likely to lead to a reward. **A**, proportion of trials in which the subjects chose the optimal action, i.e. the one more likely to be rewarded, given the brain state representing motion direction. Although on day 2 the relevant brain state was still ‘hidden’ (unreflected by the visual stimulus), subjects showed significant learning nonetheless. **B**, perceptual (state) discrimination accuracy, as leftward vs rightward motion discrimination. The correctness of the response was based on the output of the decoder. The multivariate internal signal was effectively unconscious on both days 1 and 2 (chance level accuracy) **C**, across-subject correlation between sum of rewards obtained on day 1 and 2, and individual metacognitive ability (i.e. how well one’s confidence tracks visual discrimination accuracy, computed with independent behavioural data from the decoder construction session prior to Day 1; see supplementary methods). **D**, proportion of optimal actions plotted by confidence level. From day 2, confidence (in the visual discrimination task) became predictive of selection of optimal action. Colored dots (light/dark green) represent single subjects, blue bars the mean, error bars the s.e.m. † p<0.08, * p<0.05, *** p<0.005.

During these two days, subjects reported several strategies on their action selection, e.g., looking for patterns in the random dots. Nevertheless, with time they increasingly reported pairing action selection with leftward or rightward motion discrimination responses. To note, reward contingencies were defined based on the online decoder output, not discrimination choices. On the third day, visual stimuli explicitly carried direction information inferred from brain activity by the decoder (closed-loop feedback, see figure 1 and supplementary methods). The correct state could now be easily reported (discrimination of left/right motion, mean ± s.e.m. 0.90 ± 0.01 with chance level 0.5, figure 2B) and most subjects learned to select the optimal action (N = 18, 17 showing p(optimal action) higher than chance, binomial test P(X=17|N) < 10^−3^; one-sided t-test, chance level 0.5, 0.712 ± 0.021 [t_17_ = 10.02, *P* < 10^−7^], figure 2A). Subjects also consciously reported the rule; e.g., state_left_ → action B, state_right_ → action *A*: selecting B when motion was left, and A when motion was right (binomial test *P*(X=16|N) < 10^−3^).

As a basic control test, we looked at whether an artificial neural network (ANN) trained on subjects’ multivoxel patterns could learn to choose the correct action (see supplementary methods). To avoid a trivial formulation, the ANN was trained either with subjects’ perceptual choices and several optimization runs; or with the real optimal action labels but with stochastic gradient descent on a single or few training run. Results from these simulations indicate that ANNs have difficulty solving even a small problem (e.g., using pre-selected voxels) if they are not allowed longer time-scales (several sweeps through the trials) for training and optimization (see supplementary figure 2).

The above-chance gambling performance shown by subjects from the early stages indicate that RL happened to some degree. RL could have resulted from two (non-exclusive) processes: 1) a state-dependent RL process where the update rule depends on both state (decoded motion direction) and action (eq. 1 in supplementary methods); 2) a state-free RL process where the agent simply selects the action associated with the highest expected value (regardless of the state, eq. 2 in supplementary methods). The state-dependent model assumes that the agent performs an active inference / estimation of the unconscious brain state. The state-free model conversely is a relatively naive process, in which the agent merely tries to maximize gains considering the action outcomes, and would just follow any average bias of the multivoxel patterns to left or right motion representation. However, computational modelling analyses utilizing state-dependent and state-free variants of a standard RL algorithm (9), suggest that state-dependent RL is better in capturing subjects behaviours. Increases in optimal action selection probabilities between day 1 and day 2 significantly correlated only with improved fits of the state-dependent RL model (Pearson r = −0.73, *P* = 0.0007, supplementary figure 3A). Moreover, by using the individual estimated learning rates we computed the empirical contribution to above-chance gambling performance by each RL process (state-dependent vs. state-free). Results indicate that from day 2 optimal action selection was largely driven by a state-dependent RL policy (supplementary figure 3B). Thus, we can conclude that the brain can estimate its relevant but unconscious state and utilize it in RL to attain above-chance optimal action selection. But, this is a computationally formidable problem to search the low-dimensional state among very high-dimensional unconscious brain dynamics only by trial and error. How can this curse of dimensionality be resolved?

The conceptual model introduced earlier (6) postulates that metacognition is instrumental for the rapid discovery of relevant RL states. Specifically, confidence in the perceptual discrimination could reflect the degree to which unconscious brain states are uncovered. Furthermore, confidence has been previously associated with RL in the context of perceptual decisions (10, 11). Hence, we hypothesized that there would be a correlation between confidence and RL measures, increasing across days even before the relevant motion information was explicitly presented.

To test this, we quantified metacognitive ability, computed as meta-d’ (*12*) and independently assessed on data from the previous decoder construction stage (see supplementary methods). Roughly, this measures the trial by trial correspondence between confidence judgements and the accuracy in perceptual choices. We hypothesized (*6*) that this could predict RL performance over sessions. Indeed, we found that meta-d’ correlated with the normalized sum of rewards obtained in the first two days when no coherent motion was present in the dot motion stimuli (permutation test, Pearson r = 0.48, *P* = 0.02, figure 2C). Furthermore, the probabilities of optimal action selection increased with higher decision confidence (linear mixed-effects [LME] model, data from all days, interaction between fixed effects ‘day’ and ‘confidence’ β = 0.041, *P* = 0.0017; data restricted to day 2, ANOVA marginal tests factor ‘confidence’: F1,62 = 8.5, *P* = 0.0049, day 3: F1,68 = 31.05, *P* < 10^−5^, figure 2D). This result was further supported by confidence differences in perceptual discrimination and to the extent that subject-level strength of confidence being predictive of action selection correlated with that of perceptual discrimination (see supplementary figure 4).

One possible concern is that this pattern of findings may simply arise randomly. However, a yoked control experiment in which new naive subjects received trial sequences from the main experiment did not reproduce the same results (supplementary figure 5).

Beyond these findings, we assessed the effect of metacognition on state-dependent RL (eq. 1 in supplementary methods) in greater detail with further computational analyses. To this end, we estimated trial-by-trial reward prediction error (RPE), which reflects the degree of learning in the gambling task. To note, the main assumption for this analysis is that the RL process (at least until day 3) is unconscious. But if the brain has to learn some form of mapping between states (patterns of activity) and actions, then it should also store an approximation of the expected value of RL state-actions pairs (defined by the decoder output [state], and action [choosing A or B]). Therefore, we computed RPE explicitly, as if the unconscious RL processes were already set up to estimate action values separately depending on unconscious motion direction. The trial-by-trial magnitude of RPE (unsigned RPE [|RPE|]) was log-transformed and binned by the confidence level reported on each trial (figure 3); data were then analysed with LME models (full LME models’ specification in supplementary methods). Significant coupling between |RPE| and confidence emerged from day 2, with high confidence associated with low |RPE| and vice-versa low confidence with higher |RPE| (data from all days, significant interactions between fixed effects ‘day’ and ‘confidence’: β = −0.054, *P* = 0.002; data restricted to day 2, ANOVA marginal tests factor ‘confidence’: F1,2312 = 4.01, *P* = 0. 045). This effect of confidence emerged in parallel with above chance gambling performance (see figure 2A) and confidence becoming predictive of optimal action selection (see figure 2D). As with action selection, on day 3 the effect of confidence on RPE was at its strongest (data restricted to day 3, ANOVA marginal tests factor ‘confidence’: F1,2241 = 50.63, *P* < 10^−10^). In terms of RL, the problem in this task is not for an agent to figure out directly the association *per se* (which would be rather trivial), but rather have a closer estimate of the real RL state *itself* (defined by a multivoxel pattern of activity). We interpret these results as indicating that metacognition may be able to access richer unconscious information about the RL state, because it is needed for reward accumulation.

**Figure 3:**
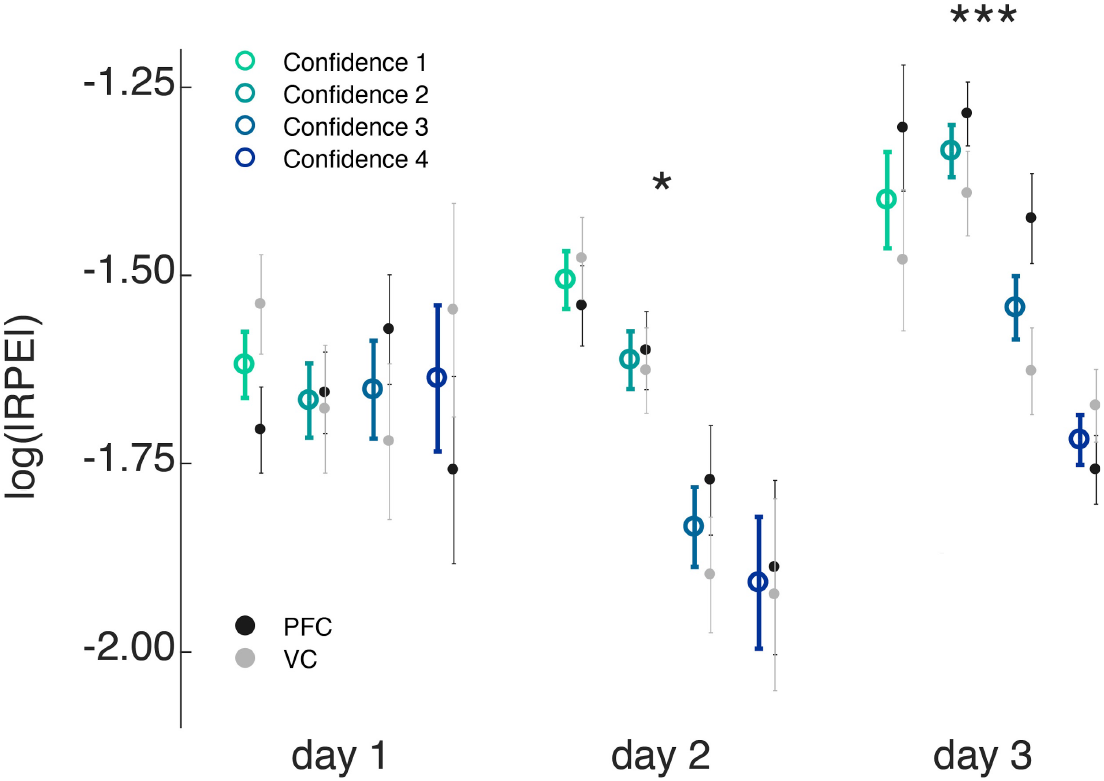
Computational modelling of behaviour: metacognition helps fast state-dependent reinforcement learning (RL). We computed reward prediction error (RPE) based on the state-dependent version of a standard RL algorithm (*9) Q*(*s,a*) ← *Q*(*s,a*) + *α* (*r* – *Q*(*s,a*)), which reflects the degree of learning in the reward action selection task. In this equation, s and a represent the decoder output (state) and the subject’s action, respectively. |RPE| data were log-transformed prior to LME model(s) fitting. The magnitude of RPE was modulated by confidence from day 2: Higher confidence in the visual discrimination task was associated with smaller absolute RPE, meaning that a high confidence choice has lower probability to result in an unexpected outcome. Coloured circles represent the mean across all subjects pooled, light/dark grey circles represent the mean across subjects pooled from VC and PFC groups, respectively; error bars the s.e.m. * p<0.05, *** p<0.005

## Neural mechanisms

At the onset of RL, cortico-basal ganglia loops are predicted to be uniformly activated in a parallel search for the relevant (unconscious) states (*6, 13*), alongside the basal ganglia (*14*). Thus, we should observe RL (RPE) -related activity in many areas of the cerebral cortex. As RL progresses, automatic selection of few, relevant loops should progress too. Because neural activity within these loops reflects the unconscious RL state, RPE-related activity in cerebral cortex and basal ganglia will shrink to few, concentrated, regional hubs. Supporting this view, recent evidence has shown that RPE correlates dynamically change over time (*13, 15*). The present results indicate that through RPE, the brain undergoes a global search initially spanning occipital and parietal cortices, anterior insula, several PFC subregions as well as anterior cingulate cortex, basal ganglia (day 2, figure 4A), that eventually converges to reward and belief processing hubs such as the basal ganglia, posterior parietal cortex and anterior lateral prefrontal cortex (day 3, figure 4A). Activity in the anterior cingulate cortex on day 2 is likely linked to the intensive action-selection search and model updating underpinning learning (*16*).

**Figure 4:**
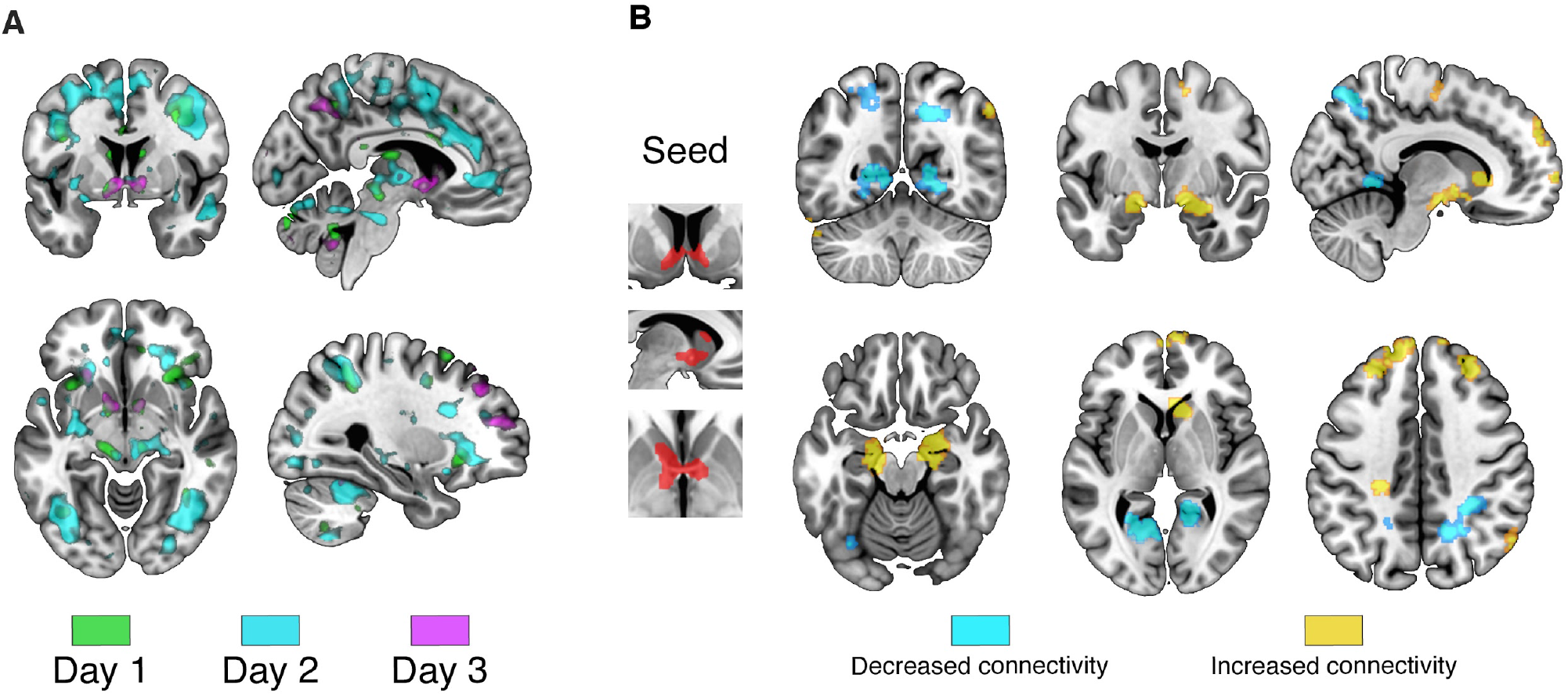
Neural correlates of parallel RL-state search. **A**, RPE correlates across the whole brain. Statistical parametric maps were generated with a general linear model with RPE as parametric regressor. View: × = −6 (top right), 28 (bottom right), y = 2, z = −8. Maps plotted at p < 0.005 (t ≷ 2.9, uncorrected). **B**, functional connectivity analysis. The seed region in the basal ganglia was defined from the RPE analysis of day 3 – independent data, collected after the last resting-state scan. View: x = 11, y = −58 (top left), −5 (top centre), z = −18 (bottom left), 2 (bottom centre), 40 (bottom right). Statistical parametric maps plotted at p < 0.005 (t ≷ 2.9, uncorrected) and cluster threshold k > 30. Maps were created by applying an F-test with contrast [−1 0 1] over the 3 resting-state scans, one-sided to test for increases (yellow) or decreases (light blue) in connectivity.

Recent co-activation of two brain areas and acquisition of knowledge or skills are believed to change resting state functional connectivity (*17–19*). Given that in our experimental setup, RL states are defined by a unique set of voxels in a predefined region (VC or PFC), one likely effect of learning is the strengthening of connections between specific brain regions and the basal ganglia that encodes and learns RPEs. Resting-state scans were collected each day, prior to the learning task (see supplementary methods); the seed region for the analysis was defined by the voxels in the basal ganglia found to be significantly correlated with RPE on day 3 (data independent of all resting-state scans, small inset in figure 4B). Clusters of voxels in the dlPFC and frontal poles showed increasingly higher correlation with activity fluctuations in the seed basal ganglia region after the first two-day RL sessions (figure 4B). Furthermore, bilateral anterior hippocampus also showed increased connectivity with basal ganglia. In line with these results, the above-mentioned regions have been recently linked to the construction of abstract representations (*20, 21*). Increased connectivity was found within basal ganglia as well as with thalamus and cerebellum subregions (figure 4, supplementary figure 6) possibly reflecting the autonomic nature of the learning process. Finally, notwithstanding the fact that each subject had a unique set of voxels utilized for the online decoding, subtle group-specific changes in functional connectivity could be detected (supplementary figure 6). Connectivity between basal ganglia and dlPFC clusters were enhanced in the dlPFC group, while the connectivity between basal ganglia and a cluster in the occipital cortex was strengthened in the VC group.

Results have so far indicated that metacognition interacts with RL, and that PFC and basal ganglia could be a potential neural substrate for this interaction. The metacognitive process over unconscious representations could use RPE to evaluate how close an estimated RL state is to a real RL state (6). From this viewpoint, if learning progresses, neural representations of confidence, RL state and RPE should become more synchronized.

We found that confidence ratings correlated with the trial-by-trial fMRI multi-voxel distance of the brain state from the decoder boundary defining the RL task states (figure 5A). Importantly, this correlation measure increased towards the end of random dot presentation, before perceptual decisions. Confidence seems to track the amount of information available in favour of one or the other state: the further the distance from the decoder’s decision boundary, the more the evidence for a given state. Metacognition essentially could provide a means of accessing the lowdimensional manifold where task goals are defined.

**Figure 5:**
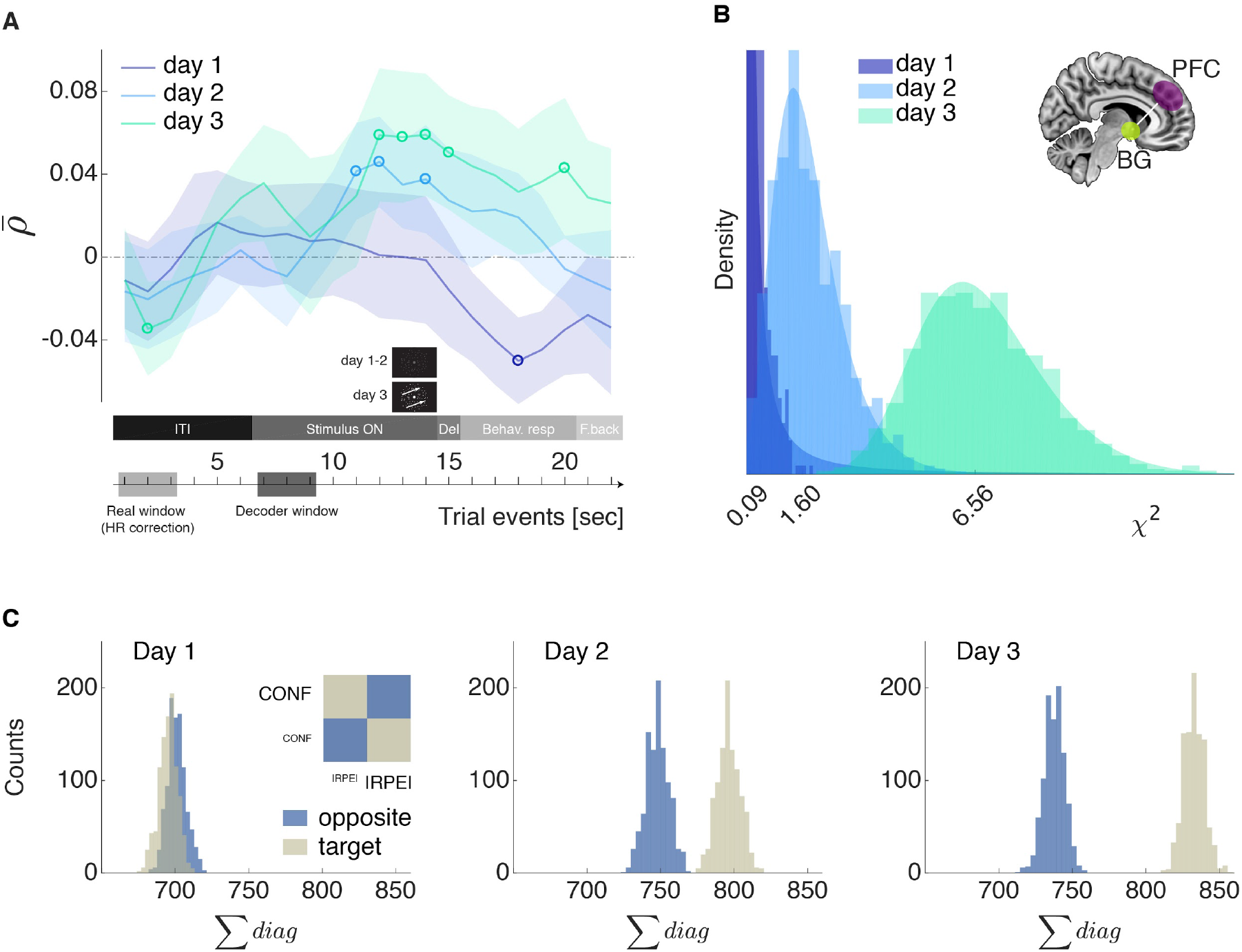
Synchronization between confidence, unconscious state and reward-prediction error. **A**, confidence judgements correlate with the activation patterns of the unique voxels used by the decoder to define the RL states. That is, reflecting the role of metacognition in the learning progression, confidence correlates with the implicit amount of RL-state evidence. Data points were shifted leftward by 6 sec to account for hemodynamic delay. Spearman correlation was computed for each subject between trial-by-trial confidence judgements and the TR-by-TR dot product between decoder weights and voxels activities. Correlation coefficients were Fisher-transformed to compute statistics. Y-axis: <u>*ρ*</u>, line is the group mean, shaded areas represent s.e.m., circles represent mean data points that are significantly different from zero, after correction for multiple comparisons (Holm-Bonferroni). **B**, multivoxel pattern association between basal ganglia and prefrontal cortex supporting confidence – RPE correlation. A decoder for confidence was built from multivoxel patterns in the PFC, while a decoder for |RPE| was constructed in the basal ganglia. For each day, the original data were randomly resampled 1000 times at the subject level and then pooled over the population to create *χ2* distributions (plotted as histograms and as shaded areas from a standard generalized extreme value fit) to indicate the degree of association between confidence and RPE. **C**, the histogram plots display the distributions over 1000 resampling runs (same as above), all subjects pooled, of the sum of the occurrences of predicted confidence pairing with predicted |RPE|. *Target* (gold coloured) is the sum of the occurrences where predicted high confidence paired with predicted low |RPE| (or predicted low confidence with predicted high |RPE|). *Opposite* (blue coloured) is the sum of occurrences where predicted high confidence paired with predicted high |RPE| (or predicted low confidence with predicted low |RPE|). The increased dependency between multivoxel patterns was specific such that, from the 2nd day, the *target* distribution of sums became greater than the *opposite* category.

Finally, given that at the computational level confidence and RPE become correlated with learning, their representations should follow the same course. To test this model prediction, we constructed a decoder for low vs. high confidence in the PFC, and a decoder for low vs. high |RPE| in the basal ganglia (see supplementary methods). By tabulating the outputs of the two decoders, χ^2^ statistics can be computed to quantify the degree of association between confidence and RPE. One thousand bootstrapped runs were calculated for each RL session: the distributions show a marked shift towards higher χ^2^ values from day 1 to day 3 (figure 5B). This implies that the independence of the decoders’ outputs decreased with learning. That is, since these decoders base their predictions on patterns of voxels activities, confidence and RPE representations became more coupled at the multivoxel level. This effect is specific for the pairs of interests (low confidence – high |RPE| and high confidence – low |RPE|, figure 5C).

## Discussion

Two main questions were addressed in this study: Can human subjects learn to make strategic use of high-dimensional unconscious brain states? What is the putative mechanism and neural substrate of this ability?

The novel closed-loop neurofeedback design adopted here granted a unique opportunity to demonstrate the ability of the human brain to learn to use high-dimensional unconscious representations. We show that such a problem can be learned within a limited number of samples, without explicit presentation of the relevant knowledge (not even in masked form), nor the presence of relevant prior hard-wired information. The data reported here support a possible solution implemented by the brain. We suggest that metacognition can access unconscious states and form higher-order abstract representations, when necessary to drive efficient RL. The ability to learn hidden features in high dimensional spaces is supported by an initially activated, distributed, and parallel neural circuitry that involves the basal ganglia and PFC. Such circuitry provides the neuroanatomical basis for the interaction between metacognitive and RL modules. Previous studies have highlighted the functional relevance of parallel cortico-basal loops in terms of RL and cognition (22, 23), as well as the role played by metacognition in RL (10, 24).

Understanding how the brain can access, modulate and use its own latent representations can be instrumental in devising new learning or rehabilitation protocols, even bypassing conscious strategies.

One important question concern whether metacognition is causally related to reward learning, or whether the interaction between confidence judgements and RL processes is bidirectional. Our results seem to indicate that confidence has a direct role in allowing RL to operate in a reduced state-space (see figures 2C-D and 3). Yet, action outcome / RPE also influence *future* confidence ratings. Interestingly, this effect may arise earlier in the course of learning than that of confidence on RPE (see supplementary figure 7). This implies that as is the case with attention (25) and memory (26), metacognition and reinforcement learning processes probably interact repeatedly in time, with specific directionalities.

One may argue that, because the closed-loop RL task involved presentation of random dot-motion and a discrimination choice, subjects’ attention was already directed towards the unconscious state. Furthermore, if sensory inference is the result of combining what we see with priors we have for interpreting information, then it could be possible that the subjective experience when looking at random motion could have indexed the RL state. That is, since no information on the state is given, the prior could take over and more strongly influence what is subjectively perceived; since the prior also drives which “probability of reward” scenario applies, it may be a possible explanation for how humans learn to gamble and make judgements during the task. Nevertheless, these seem unlikely in the light of decoding of unconscious representations taking place *before* the beginning of random dot presentation (accounting for hemodynamic delay), and perceptual discrimination being not different from chance on both day 1 and 2. Even if there were implicit prior knowledge that a representation of motion direction is the relevant state, it would be unlikely for the brain to have priors on the *spatial localization* (PFC or VC) *and sparse selection* of about 100 voxels used in the RL sessions.

Besides the importance of demonstrating that human subjects can learn to use hidden, unconscious task-relevant information, the question asked here can be extended to consider the significant problem of dimensionality. In statistical learning theory the generalization error follows the relation *e* ∝ *d*/(2*n*)(*27*), where *d* is the number of dimensions and *n* the sample size. Successful artificial neural networks (*28–30*) decrease the effective number of dimensions through regularization and dropout, but still require exceedingly large training samples, particularly when states are hidden and uncertain. Because the stream of incoming perceptual information and the representational space itself (in the brain) are both high dimensional, in order to learn quickly the brain has to operate not at the feature level, but at a rather more abstract level (*31*). Together with metacognition, other cognitive functions such as episodic memory or attention may participate in this process: select few, relevant features to allow faster RL processes (*6, 25, 32, 33*).

Exploiting unconscious states and reducing the dimensionality of the search space should thus be intimately linked. In the brain, synchronization of neurons through electrical coupling or synchronization between brain areas via higher order cognitive functions have been proposed as neural mechanisms controlling degrees-of-freedom in learning (*6, 34, 35*). Metacognition and consciousness could thus have a clear computational role in adaptive behaviour and learning (31, 36), a point that is particularly interesting given the current success in developing artificial agents. Notably, Dehaene *et al*. (*36*) discussed these aspects precisely from the viewpoint of their significance for artificial intelligence (AI) – consciousness would allow information to be flexibly broadcasted to distant nodes while metacognition could represent error-or reality-monitoring, as well as the degree of certainty in current beliefs. Our study indicates converging computational roles: higher order, low-dimensional representations that can be flexibly used by RL.

Finally, how do these findings integrate within the bigger picture of AI and neuroscience? It is beyond the current scope to provide an explicit implementation of how metacognition and RL may interact at the neural level. Nevertheless, this is the first step in a direction we envision to be of some importance. In particular, work towards endowing artificial agents with self-monitoring capacities or the ability to operate at different representational levels (feature level, concept level, etc) may bridge the still large gap between human and AI performances in real-world scenarios, beyond pattern-recognition problems. Neuroscience-based principles such as the ones elucidated here can provide the necessary seeds to develop cognitively-inspired AI algorithms (37) and is going to be a core aspect of work in neuroscience and machine learning.

## Conflicts of interest

None

## Acknowledgements

We thank Ben Seymour, Kenji Doya, Brian Odegaard, Brian Maniscalco, Vincent Taschereau-Dumouchel, Ben Smith, Matthias Michel for helpful comments on earlier versions of the manuscript. A.C and M.K. were supported by AMED (Japan, grant number JP18dm0307008), A.C. was further supported by JST ERATO (Japan, grant number JPMJER1801), H.L. was supported by the National Institutes of Health (US, grant number R01NS088628).

## Materials and Methods

### Subjects

22 subjects (23.6 ± 4.0 y.o.; 5 females) with normal or corrected-to-normal vision participated in stage 1 (motion decoder construction). One subject was removed because of corrupted data; one subject withdrew from the experiment after stage 1. We initially selected 20 subjects, of which one was removed after the first day of RL (RL) training due to a technical issue (scanner misalignment between stage 1 and new sessions), while a second subject was removed due to a bias issue with online decoding (all outputs were of the same class). Thus, 18 subjects (23.4 ± 3.3 y.o., 5 females) attended all neurofeedback RL sessions. All results presented are from the 18 subjects that completed the whole experimental timeline, with a total of 72 scanning sessions.

All experiments and data analyses were conducted at the Advanced Telecommunications Research Institute International (ATR). The study was approved by the Institutional Review Board of ATR. All subjects gave written informed consent.

### Stage 1 (day 0): Behavioural task

The initial decoder construction took place within a single session. Subjects engaged in a simple perceptual decision making task (4): upon presentation of a random dot motion (RDM) stimulus they were asked to make a choice on the direction of motion and then rate their confidence about their decision (supplementary figure 1). The choice could be either right or left, and confidence was rated on a 4-point scale (from 1 to 4), with 1 being the lowest level – pure guess, and 4 the highest level – full certainty. The task itself was identical to that used in a previous study, see (4) for reference.

The coherence level of the RDM stimuli was defined as the percentage of dots moving in a specified direction (left or right). Half of the trials had high motion coherence (50%). The latter half had threshold coherence (between 5-10%). On those threshold trials, coherence was individually adjusted to maintain the task accuracy at perceptual threshold, ~75% correct.

The entire stage 1 session consisted of 10 blocks. A 1-minute rest period was provided between each block upon subject’s request. Each block consisted of 20 task trials, with a 6 sec fixation period before the first trial and a 6 sec delay at the end of the block (1 run = 292 sec). Throughout the task, subjects were asked to fixate on a white cross (size 0.5 deg) presented at the centre of the display. Each trial started with an RDM stimulus presented for 2 sec, followed by a delay period of 4 sec. Three sec were then allotted for behavioural responses (direction discrimination 1.5 sec, confidence rating 1.5 sec). Lastly, a trial ended with an intertrial interval (ITI) of variable length (between 3 and 6 sec); see supplementary fig. 1.

Because subjects were in the MR scanner while performing the behavioural task, they were instructed to use their dominant hand to press buttons on a diamond-shaped response pad. Concordance between responses and buttons was indicated on the display and, importantly, randomly changed across trials to avoid motor preparation confounds (i.e., associating a given response with a specific button press).

### fMRI scans: acquisition and protocol

The purpose of the fMRI scans in stage 1 was to obtain fMRI signals corresponding to viewed direction of motion (e.g., rightward and leftward motion) to compute the parameters for the decoders used in stage 2, the online RL training. All scanning sessions took place in a 3T MR scanner (Siemens, Prisma) with a 64-channel head coil in the ATR Brain Activation Imaging Centre. Gradient T2*-weighted EPI (echoplanar) functional images with blood-oxygen-level-dependent (BOLD) sensitive contrast and multi-band acceleration factor 6 were acquired. Imaging parameters: 72 contiguous slices (TR = 1 sec, TE = 30 ms, flip angle = 60 deg, voxel size = 2×2×2 mm^3^, 0 mm slice gap) oriented parallel to the AC-PC plane were acquired, covering the entire brain. T1-weighted images (MP-RAGE; 256 slices, TR = 2 s, TE = 26 ms, flip angle = 80 deg, voxel size = 1×1×1 mm^3^, 0 mm slice gap) were also acquired at the end of stage 1. The scanner was realigned to subjects’ head orientations with the same parameters on all days.

### fMRI scans: preprocessing for decoding

(BOLD) signals were thus obtained for all behavioural measures associated with the task. The fMRI data for the initial 6 sec of each run were discarded due to possible unsaturated T1 effects. The fMRI signals in native space were preprocessed in MATLAB Version 7.13 (R2011b) (MathWorks) with the mrVista software package for MATLAB (http://vistalab.stanford.edu/software/). The mrVista package uses functions from the SPM suite (SPM12, http://www.fil.ion.ucl.ac.uk/spm/). All functional images underwent 3D motion correction. No spatial or temporal smoothing was applied. Rigid-body transformations were performed to align the functional images to the structural image for each subject. A grey-matter mask was used to extract fMRI data only from grey-matter voxels for further analyses. Regions of interest (ROIs) were anatomically defined through cortical reconstruction and volumetric segmentation using the Freesurfer software, which is documented and freely available for download online (http://surfer.nmr.mgh.harvard.edu/). Furthermore, visual cortex (VC) subregions V1, V2, and V3 were also automatically defined based on a new probabilistic map atlas (38). Once ROIs were individually identified, time-courses of BOLD signal intensities were extracted from each voxel in each ROI and shifted by 6 sec to account for the hemodynamic delay using the MATLAB software. A linear trend was removed from the time-courses, and further z-score normalized for each voxel in each block to minimize baseline differences across blocks. The data samples for computing the motion (and confidence) decoders were created by averaging the BOLD signal intensities of each voxel for 6 volumes, corresponding to the 6 sec from stimulus onset to response onset (supplementary fig. 1).

### Decoding: multivoxel pattern analysis (MVPA)

All MVP analyses followed the same procedure. We used sparse logistic regression (SLR) (39), which automatically selects the most relevant voxels for the classification problem, to construct binary decoders (motion: leftward vs. rightward motion; confidence: high vs. low; |RPE|: high vs. low).

K-fold cross-validation was used for each MVPA by repeatedly subdividing the dataset into a “training set” and a “test set” in order to evaluate the predictive power of the trained (fitted) model. The number of folds was automatically adjusted between k = 9 and k = 11 in order to be a (close) divisor of the number of samples in each dataset. Furthermore, SLR classification was optimized by using an iterative approach: in each fold of the cross-validation, the feature-selection process was repeated 10 times. On each iteration, the selected features (voxels) were removed from the pattern vectors, and only features with unassigned weights were used for the next iteration. At the end of the k-fold cross-validation, the test accuracies were averaged for each iteration across folds, in order to evaluate the accuracy at each iteration. The number of iterations yielding the highest classification accuracy was then used for the final computation, using the entire dataset to train the decoder that would be used in the closed-loop RL stage. Thus, each decoder resulted in a set of weights assigned to the selected voxels; these weights can be used to classify any new data sample.

Data from stage 1 (day 0) was used to train motion decoders. Pilot analyses indicated that the highest classification accuracies in PFC were attained by using high motion coherence trials alone (100 trials, 50 samples per class). Motion decoders were constructed with fMRI data from two brain regions: prefrontal cortex (PFC) and VC. These data were time-course extracted from the 6 sec from stimulus onset to response onset. Subjects were assigned to either the VC or PFC group so as to minimize the difference in overall cross-validated decoding accuracy between the two groups (see supplementary table 1 for subject-specific subregions). The mean (± s.e.m) number of voxels available for decoding was 3222 ± 309 for VC, and 4443 ± 782 for PFC. The decoders selected on average 80 ± 15 voxels in VC, and 63 ± 18 in PFC. The cross-validated test decoding accuracy (mean ± s.e.m.) for classifying leftward vs. rightward motion was 70.44 ± 2.63 % for VC, and 65.51 ± 1.35 % for PFC (two-sample t-test, t_16_ = 1.67, *P* = 0.11).

For confidence decoders, trials from stage 1 (day 0) with threshold coherence were used (100 trials), this in order to avoid potential confounds due to large differences in stimulus intensity. Because confidence judgements were given on a scale from 1 to 4, trials were first binarized into high and low confidence ratings, as described previously (4). Confidence decoders were constructed with fMRI data from dorsolateral prefrontal cortex (dlPFC, which included the inferior frontal gyrus, middle frontal gyrus and middle frontal sulcus), and time-course extracted from the 6 sec from stimulus onset to response onset. The mean (± s.e.m) number of voxels available for decoding was 6641 ± 183, and the decoders selected on average 40 ± 8 voxels. The crossvalidated test decoding accuracy (mean ± s.e.m.) for classifying high vs. low confidence was 68.77 ± 1.53 %.

For RPE magnitude (unsigned RPE) decoders, fMRI data from stage 2 was used (see sections *Stage 2 (day 1, 2, 3): online reinforcement learning training* and *Reinforcement learning modelling* for a description on the task, timing, and computation of trial-by-trial RPE). All trials from day 3 were used and, similar to confidence decoders, trials were labelled according to a median split of the unsigned RPE. If |RPE| was larger than the median, the associated trial was labelled as high RPE, and vice versa. |RPE| decoders were constructed with fMRI data from basal ganglia (which included bilateral caudate, putamen and pallidum), and time-course extracted from the 2 sec from monetary outcome presentation. The mean (± s.e.m) number of voxels available for decoding was 3583 ± 81, and the decoders selected on average 69 ±14 voxels. The cross-validated test decoding accuracy (mean ± s.e.m.) for classifying high vs. low |RPE| was 57.34 ± 0.64 %.

### Stage 2 (day 1, 2, 3): online reinforcement learning training

Once a targeted motion decoder was constructed, subjects participated in 3 consecutive days of RL online training (Fig. 1). In the RL task, state information was directly computed from fMRI voxel activity patterns in real time. The setup allowed us to create a closed loop between (spontaneous) brain activity in specific areas and task conditions (behaviour). The loop was unknown to subjects; the only instruction they received was that they should learn to select one action among two options, in order to maximize their future reward.

On each day, subjects completed up to 12 fMRI blocks; on average (mean ± s.e.m.) 9.9 ± 0.4, 11.2 ± 0.2, and 10.5 ± 0.2 blocks on day 1, 2, and 3, respectively. Each fMRI block consisted of 12 trials (1 trial = 22 sec) preceded by a 30-sec fixation period and ending with an additional blank 6 sec (1 block = 300 sec). Furthermore, on each day, before the reinforcement task, subjects underwent an additional resting-state scan of the same duration (300 sec).

The construction of an online trial observed the following rule. After a 6 sec blank ITI (black screen), the RDM was presented for a total of 8 sec. The first 6 sec were always random (0% coherence), while on day 3 the last 2 sec of RDM had coherent *(coh)* dot motion, computed as:

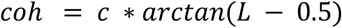

where *L* is the likelihood, the output of the motion decoder, and *c* a constant, which increased over the first half of the experimental session following a sigmoid function within the interval (0 1). Negative values indicated leftward motion, while positive values rightward motion. This allowed us to have high coherence in the latter half of day 3. Additionally, the strength of the RDM stimulus was modulated by the contrast of the dots on the black background. Contrast was set at a fixed value of 20% on day 1 and day 2 while on day 3 it increased up to 100% over the first half of the experimental session following a sigmoidal function, staying fixed thereafter. Importantly, because the operation of stimulus presentation and online decoding were performed by two parallel scripts on the same machine, the stimulus was presented in brief intervals of dot motion lasting 850 ms, followed by a short blank period of 150 ms. The presence of the blank period allowed the two processes to communicate in order to compute the new coherence level from the decoder output likelihood. Although this was effectively carried out only on day 3, the same design was used on each day for consistency between sessions. Following RDM presentation and a 1 sec blank ITI, subjects had 1.5 sec to make a state discrimination (choose leftward or rightward motion), and 1.5 sec to give a confidence judgement on their decision (on a scale from 1 to 4). Lastly, subjects had to select one of two options, A or B, in order to maximize their future reward. The reward rule for options A and B was probabilistic and determined by the decoded brain activity. Each option was thus optimal only in one state (e.g., A when left motion was decoded from multivoxel patterns, B with right motion). The probability of receiving a reward was ~80% if the choice was congruent with the rule, ~20% otherwise. A rewarded trial corresponded to a single bonus of 30Y. On each day, up to 3000 JPY could be paid to the subjects. Crucially, the reward association rule and the presence of online decoding were withheld from subjects: they were simply instructed to explore and try to learn the rule that would maximize their reward.

Because brain activity patterns *alone* were defining whether a trial was to be labelled as rightward or leftward – the experimenter had no control over the occurrence of either state (leftward or rightward motion representation). Behavioural responses could not be associated with a specific button press: pairings between buttons and responses were randomly determined on each trial and cued on the screen during response times.

### Real-time fMRI preprocessing

In each block, the initial 10 sec of fMRI data were discarded to avoid unsaturated T1 effects. First, measured whole-brain functional images underwent 3D motion correction using Turbo BrainVoyager (Brain Innovation). Second, time-courses of BOLD signal intensities were extracted from each of the voxels identified in the decoder analysis for the target ROI (either VC or PFC). Third, the time-course was detrended (removal of linear trend), and z-score normalized for each voxel using BOLD signal intensities measured up to the last time point. Fourth, the data sample to calculate the RL state and its likelihood was created by taking the BOLD signal intensities of each voxel over 3 sec (3TRs) from RDM onset. Finally, the likelihood of each motion direction being represented in the multivoxel activity pattern was calculated from the data sample using the weights of the previously constructed motion decoder. The final prediction was given by the average of the 3 likelihoods computed from the 3 data points.

### Reinforcement learning modelling

We used a standard RL model (9) to derive individual estimates of how subjects’ action-selection was dependent on past reward history tied to actions and states (state-dependent RL) or actions alone (state-free RL). State-dependent (1) and state-free (2) RL are formally described as:

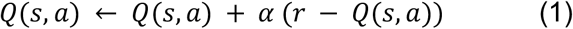

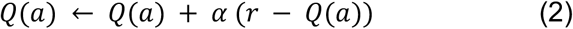

Where *Q*(*s, a*) in (1), *Q*(*a*)in (2), is the value of selecting A or B. The value of the action selected on the current trial is updated based on the difference between the expected value and the actual outcome (reward or no reward). This difference is called the reward prediction error (RPE). The degree to which this update affects the expected value depends on the learning parameter *α.* The larger *α*, the more recent outcomes will have a strong impact. On the contrary, a small *α* means recent outcomes will have little effect. Only the value of the selected action (which is state-contingent in (1)) is updated. The values of the two actions are combined to compute the probability *P* of predicting each outcome using a softmax (logistic) choice rule:

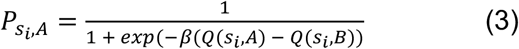

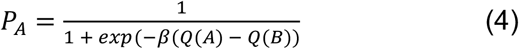

The inverse temperature *β* controls how much the difference between the two predictions values for A and B influences choices.

The two hyperparameters α and β were estimated by minimizing the negative log likelihoods of choices given the estimated probability *P* of each choice. We conducted a grid search over the parameter spaces *α* ∈ (0,1) and *β* ∈ (0,20), with 50 steps each. Rather than directly using the single point estimates, we generated the marginal likelihoods of each parameter and then used these to compute the respective expected estimates. The fitting procedure was repeated for each subject and each day (see supplementary table 2, group mean ± s.e.m). Trial-by-trial RPE measures were computed for each RL model, subject, and day, by fitting the data with the estimated parameters. RPEs were then used as inputs for offline analyses as described below.

### RPE-based analyses: parametric general linear model

Image analysis was performed with SPM12 (http://www.fil.ion.ucl.ac.uk/spm/). Raw functional images underwent realignment to the first image of each session. Structural images were reregistered to mean EPI images and segmented into grey and white matter. The segmentation parameters were then used to normalize and bias-correct the functional images. Normalized images were smoothed using a Gaussian kernel of 7 mm full-width at half-maximum.

Onset regressors beginning at the beginning of outcome presentation (reward feedback) were modulated by a parametric regressor, trial-by-trial RPE from state-dependent RL. Other regressors of no interest included onset regressors for each trial event (RDM, choice, confidence, action selection, reward outcome), motion regressors (6) and block regressors. Adding a reward regressor meant that the signal correlating with RPE was not confounded by mere reward.

Second-level group contrasts from GLM1 were calculated as one-sample *t*-tests against zero for each first-level linear contrast. Activations were reported at a cluster level threshold of k > 1000, and height threshold of *P* < 0.005 (t > 2.9). Statistical maps were projected onto a canonical MNI template with MRIcroGL (www.nitrc.org/projects/mricrogl).

### Connectivity analyses

For connectivity analyses of resting state data measured at the beginning of each session we used the CONN toolbox v.17 (www.nitrc.org/projects/conn, RRID:SCR_009550). Briefly, resting state data underwent realignment and unwarping, centred at (0,0,0) coordinates, slice-timing correction, outlier detection, smoothing and finally denoising. At the first level, we performed a seed-based correlation analysis, testing for significant correlations between voxels in a seed region and the rest of the brain. The seed was defined as the cluster of voxels within the basal ganglia that best tracked the RPE fluctuations on the last session of the RL task (day 3, independent data). The analysis was repeated for each session of resting state scanning (day 1, 2, 3). Second level group level results were calculated as one-sample t-tests against zero for each first-level contrast. We first looked at between-subjects (PFC > VC), applying between-days contrasts (day 3 > day 1) at a height threshold of *P* < 0.005 (uncorrected), and cluster size = 30. Connections to PFC and cerebellum increased over days. Conversely, the between-subjects contrast (VC > PFC) indicated increased connections between basal ganglia and cerebellum and posterior cingulate cortex (supplementary figure 4). Given these differences, we analysed separately the two groups (results reported in the main text) applying between-days contrast (day 3 > day 1) at a height threshold of *P* < 0.005 (uncorrected), and cluster size = 30. Statistical maps were projected onto a canonical MNI template with MRIcroGL.

### Statistical analyses with linear mixed effects models

All statistical analyses were performed with MATLAB Version 9.1 (R2016b) (MathWorks), both with built-in functions as well as with functions commonly available on the MathWorks online repository or custom written code. Effects of learning on behavioural data over several days and additional effects were statistically assessed using linear mixed effects (LME) models with the MATLAB function ‘fitglme’ with ‘fminunc’ as optimizer. Post-hoc tests included LME over single days, restricted to certain variables as well as two-tailed, or single-tailed where warranted, t-tests.

To evaluate the effect of confidence (levels from 1 to 4), day (1 – 3), and group (PFC, VC) on the dependent variable *y* (I: probability of selecting optimal action, II: perceptual discrimination, III: RPE from state-dependent RL), we used the general model (in Wilkinson notation): *y ~ 1 + group*day*confidence + (1\subjects)*, which included random effects (intercept) for each subject, and 8 fixed effects (intercept, group, day, confidence, group:day, group:confidence, day:confidence, group:day:confidence). Whereby a simpler model (i.e., without 3-ways interaction), *y ~ group*day + group*confidence + day*confidence + (1\subjects)* fit the data equally well (likelihood ratio [LR] test indicating no difference), results from the simpler model are reported (alongside with LR statistics). Where a significant effect of ‘day’ or interaction between fixed effects ‘day’ and ‘confidence’ and/or ‘group’ was found, post-hoc tests were carried out on data restricted to single days. For single-day data the general model *y ~ group*confidence + (1\subjects)* was used; whereby a simpler model (i.e., without interaction) fit the data equally well, results from the simpler model are reported.

The same approach was to evaluate the effect of RPE on confidence (RPE from trial-1): the same equations and procedure, just defining *y* as confidence, while RPE was treated as a fixed effect.

| l) y = probability of selecting optimal action |  |  |  |  |  |  |  |
| --- | --- | --- | --- | --- | --- | --- | --- |
| formula | $y \sim 1 + \text{group} * \text{day} + \text{group} * \text{confidence} + \text{day} * \text{confidence} + (1 \vert \text{subjects})$ | | | | | | |
| Likelihood ratio test<br>(reduced vs. full model) | LRStat = 0.089 |  | deltaDF = 1 |  | P = 0.765 |  |  |
| | $\beta$ | SE | tStat | DF | P | CI<sub>L</sub> | CI<sub>U</sub> |
| intercept | 0.588 | 0.0861 | 6.827 | 194 | $<10^{-8}$ | 0.418 | 0.757 |
| group | -0.0457 | 0.0826 | -0.553 | 194 | 0.581 | -0.209 | 0.117 |
| day | -0.0526 | 0.0374 | -1.406 | 194 | 0.161 | -0.126 | 0.0212 |
| confidence | -0.0413 | 0.0293 | -1.408 | 194 | 0.161 | -0.0991 | 0.0165 |
| group:day | -0.0278 | 0.0281 | -0.989 | 194 | 0.324 | -0.0833 | 0.0277 |
| group:confidence | 0.0312 | 0.0213 | 1.462 | 194 | 0.145 | -0.0109 | 0.0732 |
| day:confidence | 0.0409 | 0.0128 | 3.188 | 194 | **0.0017** | 0.0156 | 0.0662 |
| Day 1 [$y \sim 1 + \text{confidence} + \text{group} + (1 \vert \text{subjects})$] | | | | | | | |
| intercept | 0.491 | 0.0634 | 7.744 | 62 | $<10^{-8}$ | 0.364 | 0.617 |
| group | 0.0063 | 0.0555 | 0.114 | 62 | 0.909 | -0.105 | 0.117 |
| confidence | 0.0178 | 0.0204 | 0.87 | 62 | 0.387 | -0.023 | 0.0586 |
| Day 2 [$y \sim 1 + \text{confidence} + \text{group} + (1 \vert \text{subjects})$] | | | | | | | |
| intercept | 0.466 | 0.0475 | 9.823 | 62 | $<10^{-12}$ | 0.371 | 0.561 |
| group | -0.0236 | 0.0406 | -0.58 | 62 | 0.564 | -0.105 | 0.0576 |
| confidence | 0.0457 | 0.0157 | 2.916 | 62 | **0.0049** | 0.0144 | 0.077 |
| Day 3 [$y \sim 1 + \text{confidence} + \text{group} + (1 \vert \text{subjects})$] | | | | | | | |
| intercept | 0.381 | 0.0571 | 6.669 | 68 | $<10^{-7}$ | 0.267 | 0.494 |
| group | -0.0508 | 0.0487 | -1.042 | 68 | 0.301 | -0.148 | 0.0465 |
| confidence | 0.102 | 0.0182 | 5.572 | 68 | $<10^{-5}$ | 0.0653 | 0.138 |

| II) $y$ = probability of correct discrimination | | | | | | | |
| --- | --- | --- | --- | --- | --- | --- | --- |
| formula | $y \sim 1 + \text{group} * \text{day} + \text{group} * \text{confidence} + \text{day} * \text{confidence} + (1 \vert \text{subjects})$ | | | | | | |
| Likelihood ratio test<br>(reduced vs. full model) | | LRStat = 1.02 | | deltaDF = 1 | | $P = 0.31$ | |
| | $\beta$ | SE | tStat | DF | $P$ | CI<sub>L</sub> | CI<sub>U</sub> |
| intercept | 0.451 | 0.0967 | 4.667 | 194 | $<10^{-5}$ | 0.260 | 0.642 |
| group | 0.0182 | 0.0902 | 0.202 | 194 | 0.840 | -0.160 | 0.196 |
| day | -0.0377 | 0.0430 | -0.878 | 194 | 0.381 | -0.122 | 0.0470 |
| confidence | -0.0578 | 0.0337 | -1.717 | 194 | 0.0876 | -0.124 | 0.00861 |
| group:day | 0.0266 | 0.0323 | 0.823 | 194 | 0.411 | -0.0371 | 0.0903 |
| group:confidence | -0.0102 | 0.0244 | -0.418 | 194 | 0.676 | -0.0583 | 0.0379 |
| day:confidence | 0.0695 | 0.0147 | 4.711 | 194 | $<10^{-5}$ | 0.0404 | 0.0986 |
| Day 1 [$y \sim 1 + \text{confidence} + \text{group} + (1 \vert \text{subjects})$] | | | | | | | |
| intercept | 0.394 | 0.0631 | 6.249 | 62 | $<10^{-6}$ | 0.268 | 0.52 |
| group | 0.039 | 0.0474 | 0.824 | 62 | 0.413 | -0.0557 | 0.134 |
| confidence | 0.0299 | 0.0219 | 1.37 | 62 | 0.176 | -0.0137 | 0.0736 |
| Day 2 [$y \sim 1 + \text{confidence} + \text{group} + (1 \vert \text{subjects})$] | | | | | | | |
| intercept | 0.436 | 0.0572 | 7.633 | 62 | $<10^{-8}$ | 0.322 | 0.551 |
| group | 0.0226 | 0.0505 | 0.447 | 62 | 0.657 | -0.0785 | 0.124 |
| confidence | 0.0334 | 0.0185 | 1.802 | 62 | 0.0764* | -0.00364 | 0.0704 |
| Day 3 [$y \sim 1 + \text{confidence} + \text{group} + (1 \vert \text{subjects})$] | | | | | | | |
| intercept | 0.333 | 0.0553 | 6.024 | 68 | $<10^{-6}$ | 0.223 | 0.444 |
| group | 0.0813 | 0.0486 | 1.672 | 68 | 0.099 | -0.0157 | 0.178 |
| confidence | 0.163 | 0.0174 | 9.379 | 68 | $<10^{-12}$ | 0.128 | 0.198 |

| III) $y = \text{RPE}_{\text{sb}}$ (RPE from state-dependent RL, log-transformed) | | | | | | | |
| --- | --- | --- | --- | --- | --- | --- | --- |
| formula | $y \sim 1 + \text{group} * \text{day} + \text{group} * \text{confidence} + \text{day} * \text{confidence} + (1 \vert \text{subjects})$ | | | | | | |
| Likelihood ratio test<br>(reduced vs. full model) | LRStat = 3.563 | | deltaDF = 1 | | $P = 0.059$ | | |
| | $\beta$ | SE | tStat | DF | $P$ | CI<sub>L</sub> | CI<sub>U</sub> |
| intercept | -1.868 | 0.126 | -14.78 | 6614 | $<10^{-45}$ | -2.116 | -1.62 |
| group | 0.0745 | 0.126 | 0.592 | 6614 | 0.554 | -0.172 | 0.321 |
| day | 0.242 | 0.0527 | 4.596 | 6614 | $<10^{-5}$ | 0.139 | 0.345 |
| confidence | 0.0169 | 0.045 | 0.375 | 6614 | 0.707 | -0.0713 | 0.105 |
| group:day | -0.0433 | 0.0398 | -1.086 | 6614 | 0.277 | -0.121 | -0.0348 |
| group:confidence | 0.0044 | 0.031 | 0.141 | 6614 | 0.888 | -0.0563 | 0.0651 |
| day:confidence | -0.0543 | 0.0176 | -3.089 | 6614 | **0.002** | -0.0888 | -0.0198 |
| **Day 1** [$y \sim 1 + \text{confidence} + \text{group} + (1 \vert \text{subjects})$] | | | | | | | |
| intercept | -1.689 | 0.188 | -8.98 | 2059 | $<10^{-17}$ | -2.058 | -1.32 |
| group | 0.0342 | 0.25 | 0.137 | 2059 | 0.891 | -0.456 | 0.524 |
| confidence | 0.0127 | 0.0323 | 0.393 | 2059 | 0.695 | -0.0506 | 0.076 |
| **Day 2** [$y \sim 1 + \text{confidence} + \text{group} + (1 \vert \text{subjects})$] | | | | | | | |
| intercept | -1.513 | 0.156 | -9.733 | 2312 | $<10^{-20}$ | -1.818 | -1.209 |
| group | -0.0406 | 0.202 | -0.201 | 2312 | 0.84 | -0.436 | 0.355 |
| confidence | -0.0564 | 0.0282 | -2.002 | 2312 | **0.045** | -0.112 | -0.0012 |
| **Day 3** [$y \sim 1 + \text{confidence} + \text{group} + (1 \vert \text{subjects})$] | | | | | | | |
| intercept | -1.0622 | 0.131 | -8.107 | 2241 | $<10^{-14}$ | -1.319 | -0.805 |
| group | -0.0329 | 0.153 | -0.214 | 2241 | 0.83 | -0.334 | 0.268 |
| confidence | -0.16 | 0.0225 | -7.116 | 2241 | $<10^{-10}$ | -0.204 | -0.116 |

| IV) y = confidence, (fixed effect RPE from state-dependent RL, log-transformed) |  |  |  |  |  |  |  |
| --- | --- | --- | --- | --- | --- | --- | --- |
| formula | $y \sim 1 + group*day + group*rpe + day*rpe + (1\vert subjects)$ | | | | | | |
| Likelihood ratio test<br>(reduced vs. full model) |  | LRStat = 3.061 |  | deltaDF = 1 |  | P = 0.08 |  |
| | $\beta$ | SE | tStat | DF | P | CI<sub>L</sub> | CI<sub>U</sub> |
| intercept | 1.329 | 0.119 | 11.214 | 6560 | $<10^{-27}$ | 1.097 | 1.561 |
| group | 0.549 | 0.211 | 2.595 | 6560 | **0.0095** | 0.134 | 0.963 |
| day | 0.498 | 0.0257 | 19.424 | 6560 | $<10^{-80}$ | 0.448 | 0.549 |
| rpe | 0.157 | 0.0288 | 5.453 | 6560 | $<10^{-6}$ | 0.101 | 0.214 |
| group:day | -0.157 | 0.0309 | -5.079 | 6560 | $<10^{-5}$ | -0.217 | -0.0963 |
| group:rpe | -0.0139 | 0.0209 | -0.665 | 6560 | 0.506 | -0.0549 | 0.0271 |
| day:rpe | -0.103 | 0.0127 | -8.128 | 6560 | $<10^{-14}$ | -0.128 | -0.0781 |
| Day 1 [$y \sim 1 + rpe + group + (1\vert subjects)$] | | | | | | | |
| intercept | 1.995 | 0.14 | 14.3 | 2041 | $<10^{-42}$ | 1.722 | 2.269 |
| group | 0.317 | 0.261 | 1.211 | 2041 | 0.226 | -0.196 | 0.829 |
| rpe | 0.0447 | 0.015 | 2.975 | 2041 | **0.003** | 0.0152 | 0.0742 |
| Day 2 [$y \sim 1 + rpe + group + (1\vert subjects)$] | | | | | | | |
| intercept | 2.052 | 0.149 | 13.751 | 2294 | $<10^{-39}$ | 1.76 | 2.345 |
| group | 0.297 | 0.279 | 1.066 | 2294 | 0.286 | -0.25 | 0.844 |
| rpe | -0.0291 | 0.0153 | -1.906 | 2294 | 0.0568* | -0.0591 | 0.0008 |
| Day 3 [$y \sim 1 + rpe + group + (1\vert subjects)$] | | | | | | | |
| intercept | 3.021 | 0.0993 | 30.419 | 2223 | $<10^{-168}$ | 2.823 | 3.216 |
| group | 0.0738 | 0.179 | 0.413 | 2223 | 0.68 | -0.277 | 0.424 |
| rpe | -0.125 | 0.0194 | -6.405 | 2223 | $<10^{-8}$ | -0.163 | -0.0864 |

### Artificial Neural Network modelling (multilayer perceptron)

The goal of the ANN was to solve the gambling task – that is, to correctly classify trials into left or right states (since each state was associated with one constant optimal action: finding the correct state constrains the rule to a trivial combination). We utilized a multilayer perceptron (MLP; input, hidden, output layers), fully connected, feedforward, with sigmoid activation function. The network had as many units in the input layer as there were voxels in the multivoxel patterns (1:1 mapping). The hidden layer was composed of 10 units, while the output layer had 2 units. Connection weights were randomly initialized. The training procedure was based on backpropagation with gradient descent, momentum and fixed learning rate, estimated for each individual subject from their RL modelling. The data for training and testing were the trial-by-trial averaged fMRI signals from the first 3 sec from RDM onset in the online behavioural task. The dimensionality was determined by the number of voxels: either the pre-selected voxels (s-vox) or all the voxels within a target region (a-vox). To prevent a trivial formulation which would not be much comparable to human learning, we trained the MLP in three ways:

1) as many runs as there were trials, allowing the number of available trials on each run as n-5 to n (5 samples, moving window), n being the current run, using subjects’ perceptual discrimination as labels (2 cases, s-vox and a-vox). The simulation was run once for each subject.
2) a single run, using all trials at once and optimal action labels (2 cases, s-vox and a-vox). Because the number of samples in the two classes was unequal, the sample set was randomly down-sampled.
3) over five consecutive runs, using all trials at once and optimal action labels (2 cases, s-vox and a-vox). Because the number of samples in the two classes was unequal, the sample set was randomly down-sampled.

These procedures were repeated, separately, for data from each day (day 1, 2, 3). To note, no cross-validation procedure was implemented in order to test the hypothesis that even under very relaxed conditions this sort of ANN have difficulty to solve such problem formulation.

### Offline multivoxel pattern analyses (figure 5B, C)

For each day of the RL task, we used the set of voxels selected by confidence (dlPFC) and |RPE| (basal ganglia) decoders (described in the *Decoding: multivoxel pattern analysis* section) to compute the degree of association between confidence and |RPE| at the multivoxel pattern level. For |RPE|, the data set was composed of the predicted labels (high, low |RPE|) of all trials within a day. To issue these predicted labels, we inputted the preprocessed voxel activities during the 2 TRs corresponding to action selection outcome to the |RPE| decoder. For confidence, the prediction was extended to several time points. Specifically, the search was extended to TRs 8-17 (TRs corresponding to stimulus presentation, as well as those showing high correlation between confidence and RL-state on day 2, 3). Within the range 7-15 TRs we took the averaged raw voxel activities over 3 TRs for better S/N ratio before inputting data to the confidence decoders. As such, we obtained 8 predictions for each trial, and selected the single one leading to the highest association strength between confidence and |RPE| predictions over all trials, at the subject level. Finally, we obtained two vectors of the same length (number of trials within a day) of predicted |RPE| (high, low) and confidence (high, low). These vectors from each subject were concatenated and the final degree of association was thus computed through *χ^2^* statistic. The process was repeated over 1000 resampling runs by changing the subset of trials used to compute the confidence predictions at the subject level. This allowed us to create a distribution of 1000 *χ*^2^values reflecting the overall degree of association between multivoxel patterns predicting confidence in the dlPFC and |RPE| in the basal ganglia.

At the single trial level, predicted data points were categorized according to the following labels: *target* if the prediction were high confidence – low |RPE| or low confidence – high |RPE|, and *opposite* if the predictions were high confidence – high |RPE| or low confidence – low |RPE|. For each resampling run we summed all occurrences of target and opposite, creating a distribution of 1000 values. Overlapping distributions means that there is no association.

## Supplementary figures and tables

**Supplementary figure 1.**
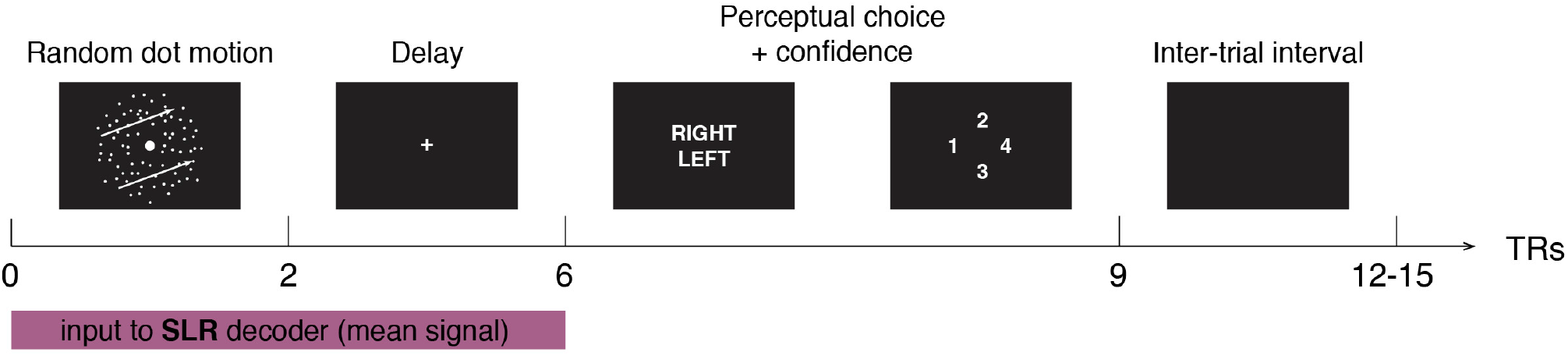
Stage 1 (day 0) behavioural task. To construct motion decoders used in stage 2 for the online training, subjects engaged in a two-choice direction discrimination task with confidence judgement while in the MR scanner. Each trial featured a random dot motion stimulus with either high or low motion coherence for 2 sec, followed by a delay period of 4 sec. Subjects were then instructed to choose a motion direction (left or right), and indicate their confidence in their choice (1 to 4) by pressing a button on a response-pad according to the positional cue presented on the screen. A trial ended with an ITI of variable length (3 to 6 seconds).

**Supplementary figure 2.**
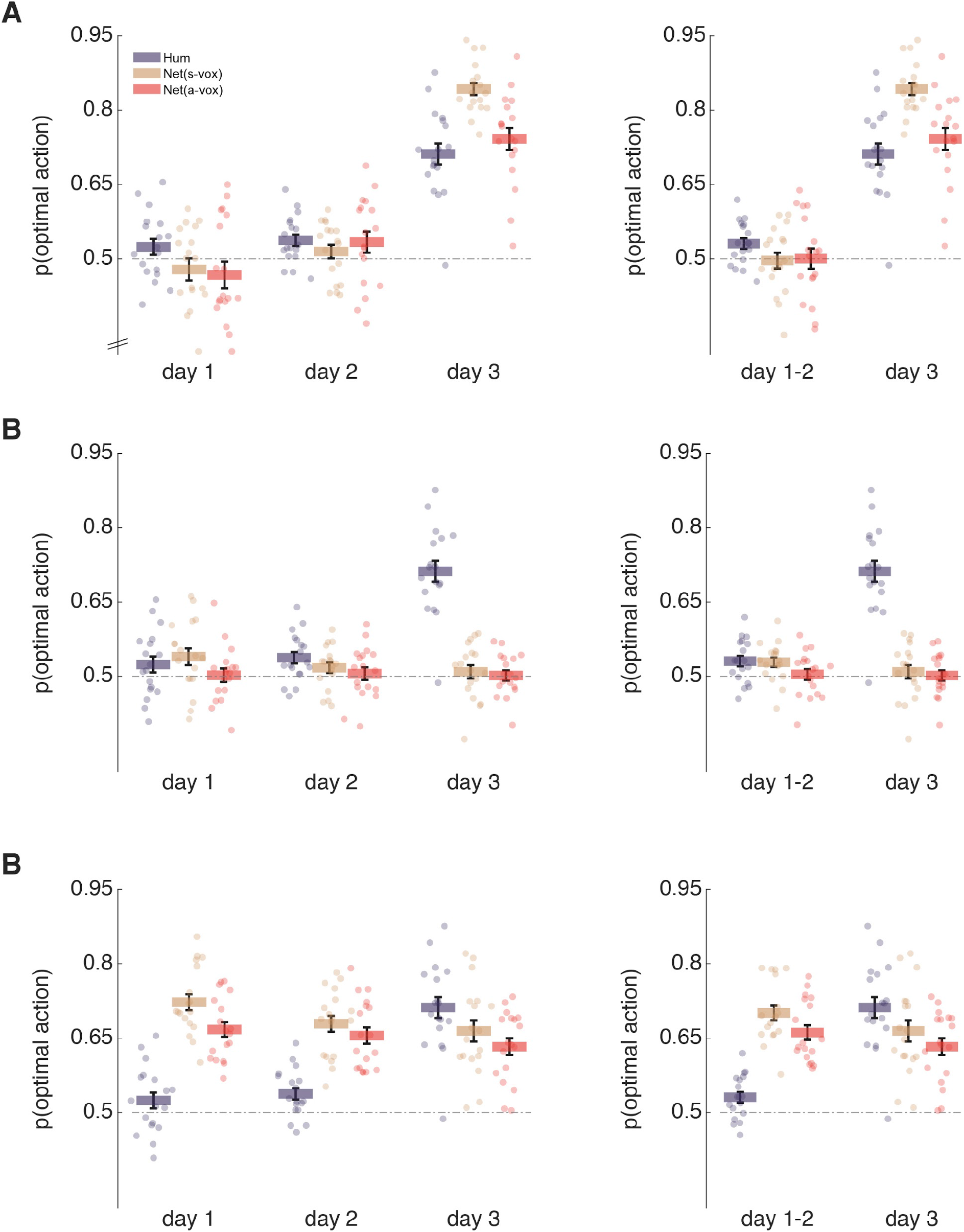
Gambling performance of multilayer perceptron. The simulation was repeated once for each subject’s data configuration, producing the same number of “behavioural” data as in the original subjects’ case. **A**, the perceptron took as input multivoxel patterns (Net(s-vox): the pre-selected voxels utilized in the online task, Net(a-vox): all voxels within the target region, either VC or PFC); the training labels were subjects’ perceptual choices, and target outputs were optimal actions. Note that when perceptual choices become accurate on day 3, the network can learn the correct association better than humans only in the very simple case of inputs consisting of pre-selected voxels. **B**, the perceptron took as input multivoxel patterns; both training labels and target output were the optimal actions. In this setting the network was trained with stochastic gradient descent, using all trials, on a single pass. Note that in this more stringent setting even providing as input the preselected voxel pattern did not result in high performance. **C**, the perceptron took as input multivoxel patterns; as in B, both training labels and target output were the optimal actions. In this setting the network was trained with stochastic gradient descent, using all trials, on five consecutive passes (during which connection weights were continuously updated). Note that with this very lenient setting the network outperformed the human on day 1 and 2, but not on day 3. In both B and C, for all networks, the learning rate was taken to be the same estimated for each subject, on each day, in the RL modelling (see supplementary methods and supplementary table 2).

**Supplementary figure 3.**
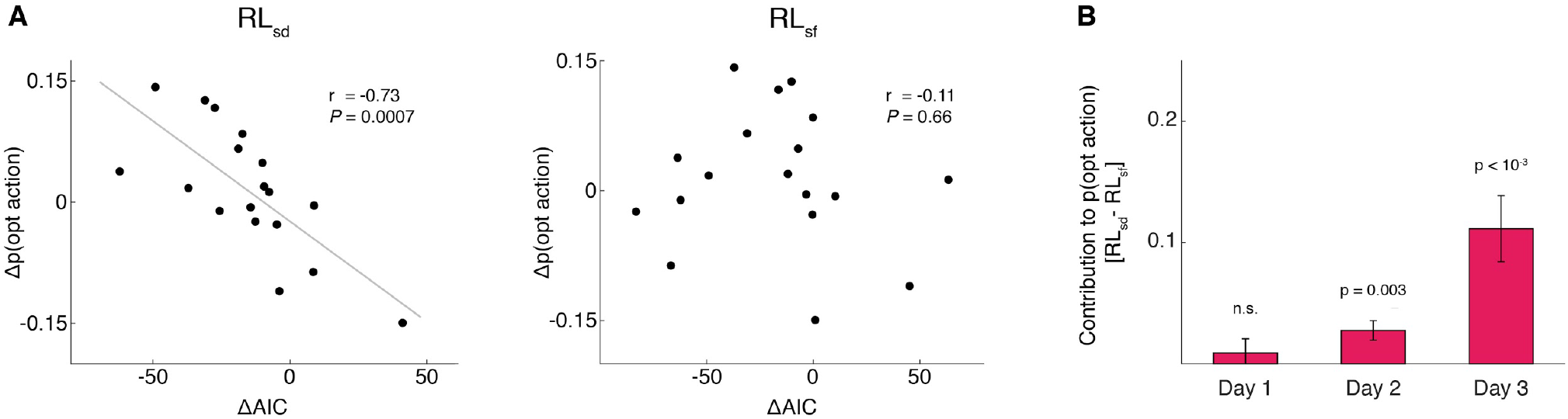
Computational modelling of action selection with reinforcement learning. Two versions of the Q-learning RL algorithm were fitted to the data. QL_sd_: state-dependent RL model based on both actions (subjects’ A/B choices) and states (decoder output), expressed as *Q*(*s,a*) ← *Q*(*s,a*) + *α* · (*r – Q*(*s,a*)); QL_sf_: state-free RL model that simply maximises actions (regardless of the state), expressed as s *Q*(*a*) ← *Q*(*a*) + *α* · (*r – Q*(*a*)). **A**, Correlation between the difference in goodness of fit computed as the ΔAIC (Akaike Information Criteria) (*40*) and changes in the optimality of action-selection for the state-dependent (RLsd) model (left) and state-free (RLsf) model (right). Differences were computed between day 2 and day 1, when the experimental settings were identical on both sessions. More negative values of ΔAIC indicate improved fits, while more positive values of Δp indicate higher optimality in action selection. **B**, Difference in the empirical contribution to subjects’ optimal action-selection by each process (state-dependent vs. state-free). Briefly, we used the maximum possible degree of optimal action selection attainable by each model: 1.0 for state-dependent, and the ratio of L/R states for state-free. The contribution of each model was then computed by normalizing the real optimal action selection (above 0.5) by the learning rates and the maximum optimal action selection attainable by each model. Positive values indicate a prevalence of RLsd over RLsf.

**Supplementary figure 4.**
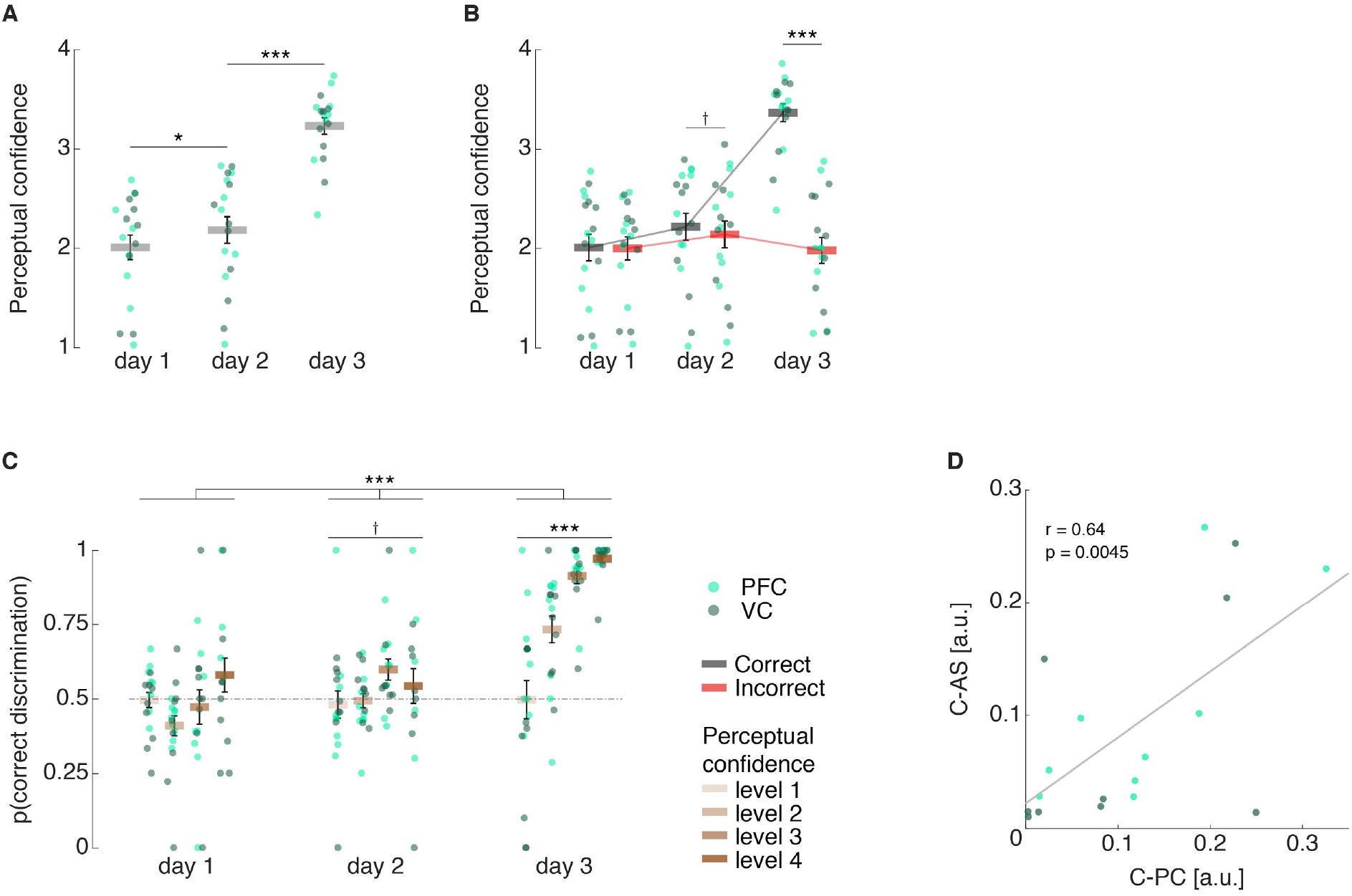
Analysis of behaviour: confidence ratings. **A**, confidence judgements for each day. Confidence increased from day 1 to day 2 and from day 2 to day 3. **B**, confidence judgements for correct and incorrect trials (discrimination, based on decoder output). From day 2, confidence was higher for correct than incorrect trials, as a form of emerging metacognitive access. On the day 3 the difference was very large, as expected in a standard decision-making task. **C**, perceptual (state) discrimination accuracy subdivided by confidence level. Data analysed with an LME model (see supplementary methods). From day 2, a weak relationship between state discrimination and confidence arose. On day 3 confidence became highly predictive of state discrimination accuracy. **D**, overall strength of confidence effect with action selection and perceptual discrimination were correlated on day 2. C-PC: measure of confidence effect on perceptual choice; C-AS: measure of confidence effect on action selection. The measure was computed as the magnitude of the averaged (signed) differences in performance over confidence levels. † p < 0.08, * p < 0.05, *** p < 0.005.

**Supplementary figure 5.**
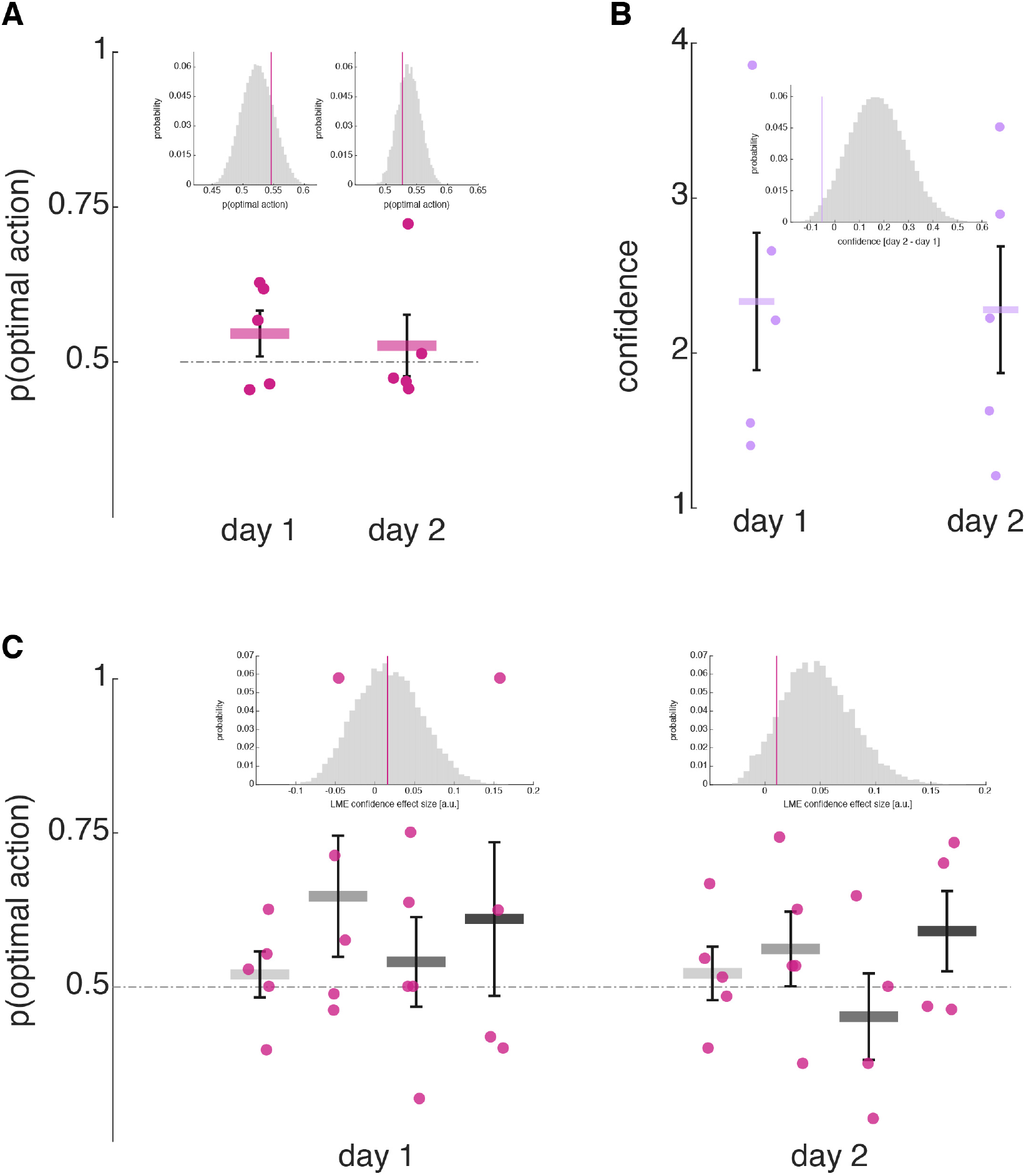
Control experiment: action selection and confidence in naive subjects without neurofeedback loop. The same task was submitted to naive subjects. As in the main experiment, physical stimuli had zero coherence. Furthermore, trials were determined exogenously: yoked trial sequences were randomly selected without replacement from those of subjects who did the full neurofeedback experiment. **A**, optimal action selection for day 1 and day 2. Inset histograms depict the distributions (day 1, day 2) of all possible random draws of N=5 from the original subjects’ data. The coloured vertical line represents the mean of the N=5 control subjects. **B**, mean confidence ratings on day 1 and day 2. No difference between the two days, as opposed to subjects from the original experiment (see supplementary figure 5A). The Inset histogram depicts the distribution of confidence difference between day 2 and day 1 of all possible random draws of N=5 from the original subjects’ data. The coloured vertical line represents the mean of the N=5 control subjects. **C**, Optimal action selection subdivided by confidence level. No effect of confidence was found [LME model, for each day parametrization as *y ~ confidence + (1 | subjects)].* AS above, the same model was run, for each day, on the original subsampled data with all possible combinations of N=5, to create a distribution of confidence coefficients for each day. On day 2 the control coefficient remains stable close to zero, but the original data distribution shows a marked shift toward higher effect sizes.

**Supplementary figure 6:**
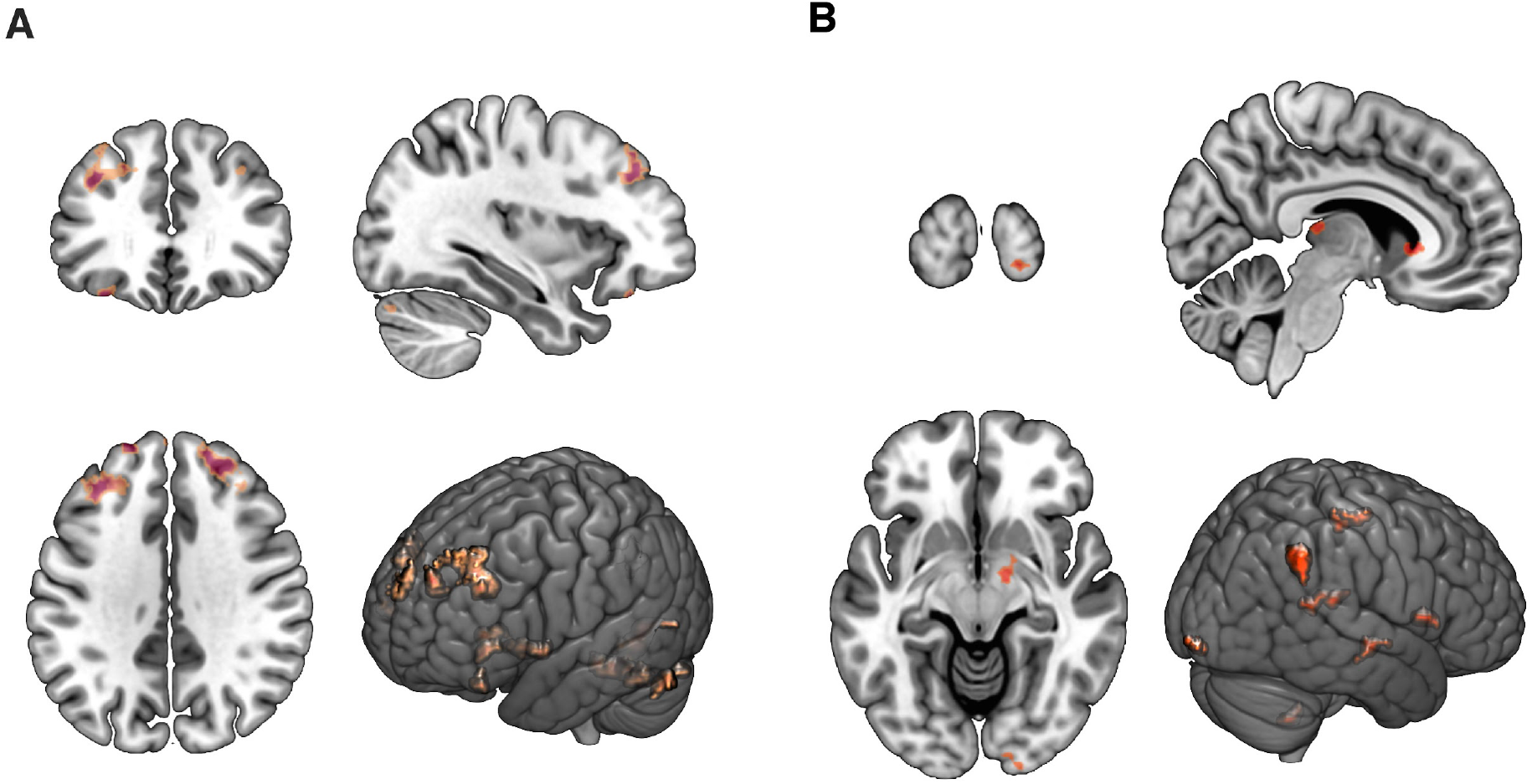
Group-specific increases in functional connectivity. The seed region in the basal ganglia was defined from the RPE analysis of day 3 – independent data, collected after the last resting-state scan. **A**, analysis restricted to subjects from the dlPFC group. View: x = −34, y = 33, z = 36. **B**, analysis restricted to subjects from the VC group. View: x = 6, y = −100, z = −10. Statistical parametric maps plotted at p < 0.005 (t > 2.9, uncorrected) and cluster threshold k > 30. Maps were created by applying a one-sided (positive) F-test with contrast [−1 0 1] over the 3 resting-state scans to test for increases in strength of functional connectivity.

**Supplementary figure 7.**
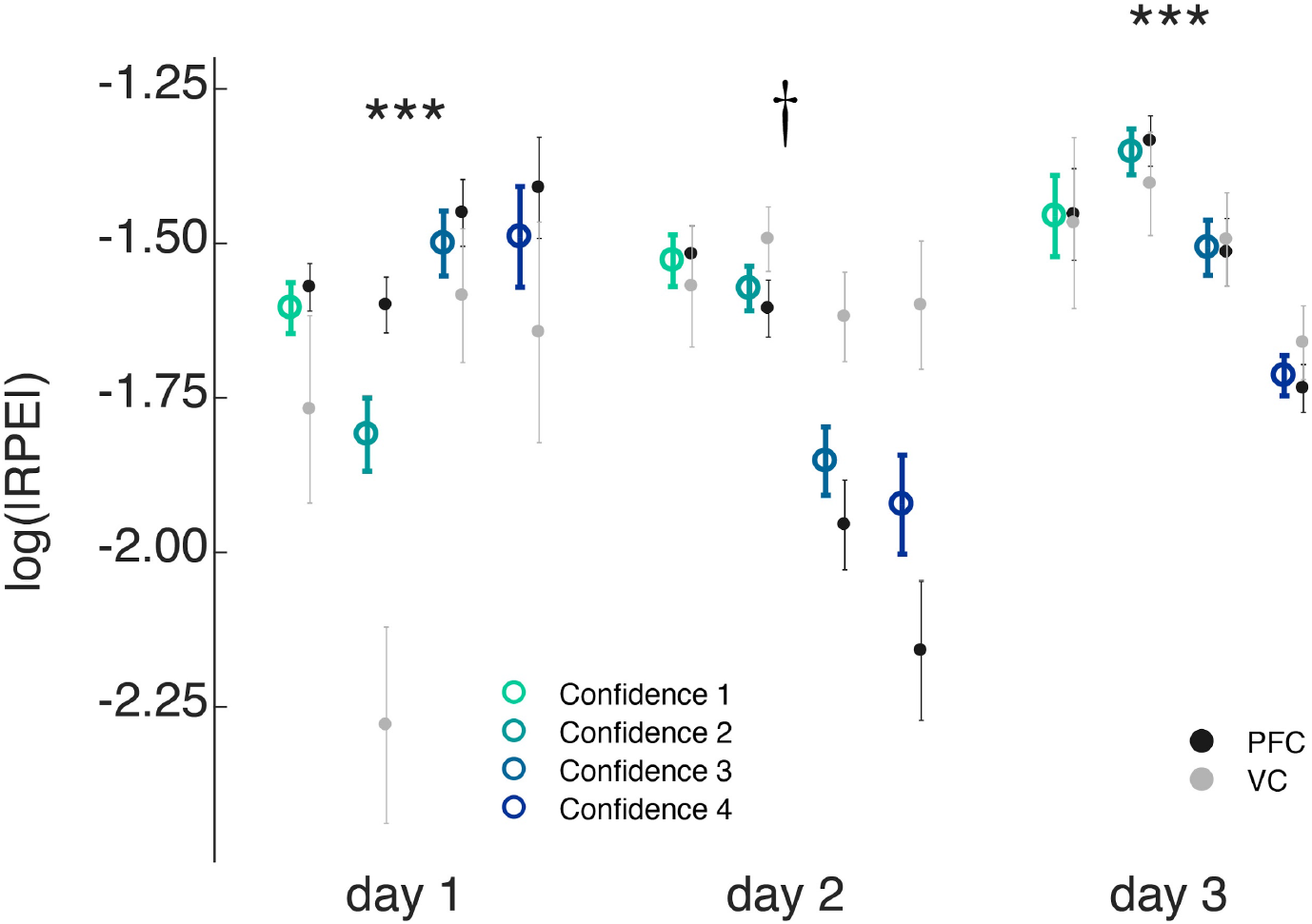
Computational modelling of behaviour: reinforcement learning effect on confidence. Reward prediction error (RPE) was computed based on the state-dependent version of a standard RL algorithm (9). The magnitude of RPE modulated confidence from the earliest stages: Smaller absolute RPE on the current trial was associated with higher confidence in the visual discrimination task in the next trial, meaning that an unexpected outcome was more likely to trigger a low confidence judgement in the next trial. |RPE| data were log-transformed prior to LME model(s) fitting. Coloured circles represent the mean across all subjects pooled, light/dark grey circles represent the mean across all subjects pooled from VC and PFC groups, respectively; error bars the s.e.m. † p<0.06, *** p<0.005

**Supplementary table 1.** Subject-specific subregions selected for motion decoder. Each subject was assigned to the PFC or VC group so as to minimize the intergroup difference in decoding accuracy (which could otherwise lead to a large confound factor). Furthermore, within each region, the decoder based on the subregion that yielded the highest mean accuracy, with the lowest difference between the two classes (i.e., leftward vs. rightward motion), was selected. The table reports for each subject the group they were assigned to (VC or PFC) as well as their individual decoder used in the stage 2 online training. PFC: prefrontal cortex, VC: visual cortex, MFG: middle frontal gyrus, MFS: middle frontal sulcus, IFS: inferior frontal cortex, dlPFC: dorsolateral prefrontal cortex (combination of IFS, MFS, and MFG), V1: area V1 of VC, V1/2: areas V1 and V2 of VC, V1/2/3: areas V1, V2, and V3 of VC.

| Subject # | Brain region | Subregion decoder |
| --- | --- | --- |
| S1 | PFC | dIPFC |
| S2 | VC | V1/2 |
| S3 | PFC | dIPFC |
| S4 | VC | V1/2 |
| S5 | PFC | MFG |
| S6 | VC | V1 |
| S7 | VC | V1/2/3 |
| S8 | PFC | MFS |
| S9 | PFC | MFG |
| S10 | VC | V1/2 |
| S11 | PFC | IFS |
| S12 | VC | V1/2 |
| S13 | PFC | dIPFC |
| S14 | VC | V1/2 |
| S15 | VC | V1/2/3 |
| S16 | PFC | MFG |
| S17 | VC | V1/2/3 |
| S18 | PFC | MFG |

**Supplementary table 2.** Estimated hyperparameters for state-dependent RL and state-free RL. Estimated values represent group mean ± s.e.m. The models were fitted on individual data, on each session (day 1, 2, 3).

|  |  | Day 1 | Day 2 | Day 3 |
| --- | --- | --- | --- | --- |
| State-dependent RL | $\alpha$ | $0.21 \pm 0.02$ | $0.19 \pm 0.02$ | $0.19 \pm 0.01$ |
| | $\beta$ | $16.51 \pm 0.25$ | $16.05 \pm 0.32$ | $16.30 \pm 0.26$ |
| state-free RL | $\alpha$ | $0.25 \pm 0.04$ | $0.23 \pm 0.03$ | $0.24 \pm 0.04$ |
| | $\beta$ | $3.36 \pm 0.79$ | $3.19 \pm 0.73$ | $2.75 \pm 0.44$ |

## Footnotes

1 Subjects were assigned into two groups, which differed in the brain regions targeted by their decoder: visual cortex (VC, N=9) or prefrontal cortex (PFC, N=9). For all analyses, brain region was treated as a between-subjects factor; unless this factor displayed a significant effect, results were reported without considering this factor, meaning subjects were effectively pooled into one cohort (N=18).

